# Connectivity, genetic diversity and adaptive potential of an endangered beetle: genomic evidence for veteran tree management

**DOI:** 10.64898/2026.09.28.755044

**Authors:** Rama Sarvani Krovi, Nermeen R. Amer, Alicja Wierzbicka, Radosław Plewa, Marcin Kadej, Tomasz Jaworski, Adrian Smolis, Tomasz Szmatoła, Maria Oczkowicz, Łukasz Kajtoch

## Abstract

Habitat fragmentation and destruction in old-growth forests represents a primary threat to specialist species, frequently resulting in the erosion of genetic diversity. The Hermit Beetle, *Osmoderma barnabita*, is a protected relic of primeval forests and an obligate saproxylic specialist whose larval development depends exclusively on the wood-mould of hollow trees. This study provides the first high-resolution, genome-wide assessment of *O. barnabita* across seven Polish forest complexes using double-digestion Restriction-site Associated DNA sequencing data from 49 individuals. Following rigorous quality control and reference bias assessment, we evaluate contemporary population structure. To distinguish between evolutionary processes, we isolated 2,920 putatively adaptive outliers from the neutral background, evaluating both neutral and adaptive genomic signals. The analyses revealed four distinct genetic clusters, confirming that the Polish metapopulation is severely subdivided. Crucially, the westernmost population in the Lower Silesian Forest emerged as the most divergent evolutionary unit (F_ST_ up to 0.360), suggesting potential hybrid origin or long-term isolation. We observed a striking linear correlation (R2=0.986) between neutral diversity and adaptive potential, indicating that genetic drift in small and isolated forest fragments is actively depleting the variation required for long-term persistence of this species. Our findings demonstrate significant gene flow restriction across all sites, suggesting that current conservation frameworks fail to encompass these highly differentiated lineages. To mitigate this genetic vulnerability, we recommend reclassifying these clusters as independent Management Units and Evolutionary Significant Units. Maintaining forest continuity and protecting local micro-habitats are essential to prevent further isolation and preserve the genomic integrity of this saproxylic flagship species.

## 1. Introduction

Global biodiversity faces intense pressure from habitat loss, a crisis most visible in Central European temperate forests (Wallenius et al., 2010). Habitat fragmentation triggers a decline in ecological function, primarily impacting obligate specialists tied to rare microhabitats (Seibold et al. 2015). Among these, saproxylic organisms, i.e. taxa dependent on decaying wood, serve as vital indicators of forest health (Bouget et al., 2013). Their vulnerability stems from the increasing scarcity of veteran trees, which provide essential substrates and microhabitats, such as cavities and wood-mould (Siitonen et al., 2012). Modern forestry has created a generational gap in tree cohorts; existing habitats are decaying without adequate recruitment of new veteran trees (Vuidot et al., 2011; IUCN 2025). Hermit beetles (*Osmoderma* spp.) are flagship taxa for European invertebrate conservation and are protected under the Habitats Directive (Maurizi et al., 2017). Once treated as a single species, the genus is now recognized as a complex of distinct lineages with partitioned ranges (Audisio et al., 2009; Maurizi et al., 2017; Mazzei et al., 2025). While *O. eremita* occupies Western and Northern Europe, *O. barnabita* inhabits Central and Eastern regions (Landvik et al., 2017). Both are classified as Near Threatened (IUCN 2025) due to habitat fragmentation and a strict reliance on hollow deciduous trees, typically *Tilia* spp. and *Quercus* spp. in older age classes (Oleksa et al. 2009). The species’ presence is tied to the historical continuity of landscapes containing long-standing hollow-tree habitats. It may also occur in anthropogenic tree stands (e.g., avenues), but typically only in landscapes that remain connected to old-growth deciduous forests; therefore, corridors formed by veteran trees are crucial for its dispersal (Kadej et al., 2016; Eckelt et al., 2018). Despite its range, the genomic characterization of *O. barnabita* in Central-Eastern Europe is sparse. Logistical and legal constraints on sampling protected species have historically limited molecular studies to low-resolution markers (Kajtoch et al., 2022). In addition, sampling protected species such as *O. barnabita* requires special permits under Polish and EU legislation, which often restricts both access to sites and the number of individuals that can be legally collected, further limiting the resolution of previous genetic studies. Prior data relied on 14 microsatellite loci (Oleksa et al., 2015) or partial mitochondrial COI sequences (Landvik et al., 2017). While the latter established deep geographic substructuring, mtDNA provides only a maternal perspective, failing to capture contemporary gene flow or the adaptive signals reflected in the nuclear genome. Consequently, we still do not know if the current protected populations are functionally connected or if we are witnessing the final stages of genomic collapse in isolated relicts.

To resolve these limitations, we employed double-digestion Restriction-site Associated DNA sequencing (ddRADseq) on 49 individuals of Osmoderma barnabita sampled from ten distinct sites across seven major Polish forest complexes. Our study is structured around three primary objectives designed to bridge the gap between genomic data and actionable conservation policy. First, we aim to generate the first high-resolution SNP dataset for O. barnabita and rigorously evaluate the methodological robustness of our findings. By comparing reference-based and de novo mapping pipelines, we test the hypothesis that both approaches will yield highly congruent population structures; we anticipate that the high marker density provided by ddRADseq will effectively overcome the technical biases inherent in any single assembly method. Second, we seek to delineate evolutionary units of conservation concern by uncovering fine-scale population structures. We hypothesize that geographically isolated forest complexes represent distinct evolutionary lineages characterized by restricted gene flow, necessitating their management as independent conservation units. Finally, we sought to assess the distribution of adaptive genetic variation and its correlation with neutral diversity. We hypothesized a strong positive correlation between the two, which would indicate that demographic forces, specifically genetic drift in fragmented habitats, are the primary drivers of genetic erosion.

## 2. Materials and Methods

### 2.1. Sample Collection and Population Designation

Genomic DNA was sourced from 49 adult *Osmoderma barnabita* collected across the following Polish forest complexes: Białowieża (BF) and Augustów (AF) in the Northeast; Carpathian Mountains (CM) and Holy Cross Mountains (HM) in the Southeast; and the Oder (OF), Barycz (DB), and Lower Silesian (SF) forests in the Southwest (Figure 1, Table 1). Beetles were captured using pheromone-baited barrier traps during June and July of 2023 and 2024. To minimize impact on this protected species, only a single middle leg was harvested from each individual before immediate release. Samples were grouped into seven populations based on their broad geographic distribution.

**Figure 1:**
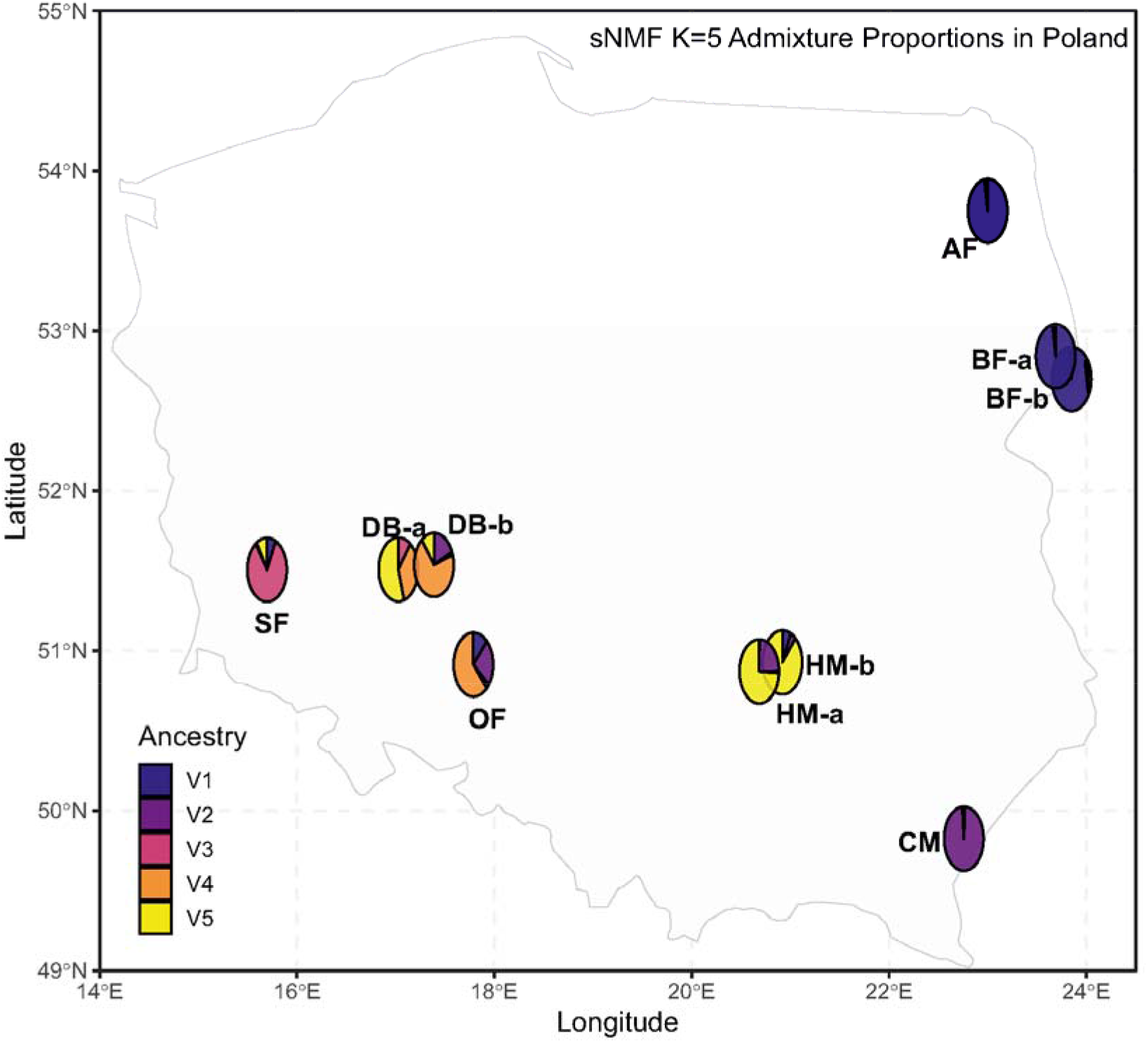
Geographic Distribution and Genetic Lineages of *Osmoderma barnabita* in nine sites across Poland. This figure illustrates the sites where *Osmoderma barnabita* samples were collected: Białowieża (BFa, BFb), Augustów (AF), Carpathian Mountains (CM), Holy Cross Mountains (HMa, HMb), Oder (OF), Barycz (DBa, DBb), and Lower Silesian (SF). It also shows the spatial admixture pie charts for K=5, which shows four geographically distinct genetic lineages across Poland.

**Table 1.** Sampling locations of *Osmoderma barnabita* in Poland.

| **Forest** | **Site** | **Latitude (N)** | **Longitude (E)** | **number of individuals** |
| --- | --- | --- | --- | --- |
| Augustów Forest | AF-a | 53.96519905 | 23.49228225 | 5 |
| Białowieża Forest | BF-a | 52.73217731 | 23.7875143 | 5 |
| Białowieża Forest | BF-b | 52.70799693 | 23.79134251 | 5 |
| Carpathians | CM-a | 49.73750168 | 22.6314045 | 4 |
| Holy Cross Mts. | HM-a | 51.05894755 | 20.70004353 | 5 |
| Holy Cross Mts. | HM-b | 50.92687448 | 20.9194108 | 5 |
| Barycz Forest | DB-a | 51.53388544 | 17.38626663 | 5 |
| Barycz Forest | DB-b | 51.47930039 | 16.91975121 | 5 |
| Oder Forest | OF-a | 50.94721397 | 17.33025884 | 5 |
| Silesia Forest | SF-a | 51.50633085 | 15.69271633 | 5 |

### 2.2. Library Preparation and Sequencing

High molecular weight DNA was extracted using the Sherlock AX kit (A&A Biotechnology). ddRAD-seq libraries were prepared following Bayona-Vásquez et al., (2019) with modifications by Amer et al., (in review). Libraries were pooled, normalized to 2 nM, and sequenced on an Illumina NovaSeq X Plus (10B, PE 150) at Admera Health (New Jersey, USA), yielding approximately 4 million reads per sample.

### 2.3. Bioinformatic Pipeline and Quality Control

Raw FASTQ data quality was assessed using FastQC and aggregated via MultiQC (Ewels et al., 2016). Raw reads were demultiplexed, cleaned, and rescued using the process_radtags module in Stacks (Catchen et al., 2013), utilizing the NheI and EcoRI restriction enzyme sites. We employed two parallel bioinformatic strategies to ensure robust results and account for potential reference bias.

1-Reference-based Calling: Reads were aligned to the *O. eremita* primary assembly (GCA_031763485.1) using minimap2 (Li, 2018). Loci were identified using the ref_map.pl pipeline.

2-De novo Calling: Genotypes were called using the denovo_map.pl and gstacks modules. We set the number of mismatches allowed between stacks within individuals to M=2 and between individuals to n=2, employing the marukilow model (var_alpha: 0.01, gt_alpha: 0.05).

Both datasets underwent rigorous filtering using the populations module and PLINK (Purcell et al., 2007). Loci were retained only if present in ≥1 population with a minimum representative rate of 80% (r=0.80). We applied a Minor Allele Frequency (MAF) ≥0.03 and a maximum missing data threshold (Geno) ≤0.30. Two Individuals with more than 50% missingness were identified and excluded from the final population maps. Locus-specific depth was verified using VCFTools (Danecek et al., 2011). Following these filters, the reference-based pipeline yielded 6150 loci and 27,364 SNPs, while the de novo approach generated 4405 loci and 9,053 SNPs. Final datasets were exported using vcfR (Knaus and Grünwald, 2017) for downstream analysis in various formats.

### 2.4. Selection Scan and Neutral Filtering

To distinguish between demographic history and selective pressures, we performed a genomic scan for outliers using pcadapt v4.3.3 (Luu et al., 2017). Using a Principal Component Analysis (PCA), we identified significant outliers (potential adaptive loci) by applying a Benjamini-Hochberg (BH) correction (FDR α=0.05). This identified 2,920 putatively adaptive loci (10.6% of the reference-based set). These were blacklisted to create a neutral-only dataset for demographic analysis and isolated as an adaptive-only dataset to evaluate evolutionary potential. The sensitivity analyses to take into consideration the differences in sample sizes and comparing neutral-only and all-loci datasets revealed no significant shifts in population clustering (PCA) and other downstream analysis. All the analysis results done using the de novo pipeline can be found in the supplementary files (Figure S1-11). Consequently, the full dataset generated using the reference based approach which consisted of 27,364 SNPs was retained for subsequent analyses. This approach ensures that the subtle signals of local adaptation are preserved alongside the broader demographic history, providing a holistic view of the genomic landscape of *O. barnabita*. (Mathur et al., 2023)

### 2.5. Population Genetic Structure and Diversity

To characterize the genetic structure of Osmoderma barnabita, we employed three complementary approaches integrating diversity metrics, kinship assessment and dimensionality reduction. All SNPs were analyzed using adegenet v2.1.10 (Jombart, 2008) and hierfstat v0.5-11 (Goudet, 2005). We calculated observed (H_o_) and expected (H_e_) heterozygosity and inbreeding coefficient (F_is_). Furthermore, to assess whether high genetic differentiation (F_st_) or low diversity might be artifacts of sampling closely related individuals (kinship), we performed a relatedness analysis. We calculated the proportion of shared alleles between all pairs of individuals using the propShared function in adegenet. This allowed us to characterize the genetic similarity across sampling sites (Figure S3). Population structure was visualized through Principal Component Analysis (PCA) to identify major axes of variation. We further estimated ancestral lineages (K) using sNMF in the LEA package (Frichot et al., 2014; Frichot & François, 2015), with the optimal K determined by the minimum cross-entropy criterion (Figure S4). Genetic subdivision was quantified using global and pairwise F_st_ based on the Weir and Cockerham (1984) estimator. Finally, we constructed a Neighbor-Joining (NJ) tree based on a bitwise genetic distance matrix (calculated via poppr) to provide a phenetic representation of the genetic proximity between populations without the requirement of a rooted phylogeny. (Figure S1, S4).

### 2.6. Genomic Erosion and Adaptive Potential

To evaluate how fragmentation impacts the species’ ability to adapt, we compared HE and Ar between the neutral and adaptive genomic compartments. We quantified genomic erosion by performing a linear regression (Pearson correlation) of adaptive HE against neutral HE across all populations. This allowed us to determine if genetic drift in isolated fragments is actively depleting the variation required for long-term persistence.

### 2.7. Estimated Effective Migration Surfaces (EEMS)

We modeled spatial gene flow using EEMS (Petkova et al., 2016) to identify migration corridors and barriers. Using 27364 filtered SNPs, we performed an MCMC analysis (250 demes) for 500,000 iterations (100,000 burn-in) (Figure S5). The study area was defined using a convex hull with a 0.5° buffer. Convergence was verified via log-posterior trace plots and the correlation between observed and fitted genetic distances.

### 2.8. COI Barcoding

To complement the ddRADseq data, we amplified a segment of the cytochrome oxidase subunit I (COI) gene using genus-specific primers (Audisio et al., 2009). All samples from SF and OF, along with random representatives from other forests, were sequenced using a BigDye Terminator v.3.1 kit on an ABI 3100 sequencer. Sequences were compared against existing *Osmoderma* records in GenBank via BLAST.

## 3. Results

### 3.1. Quality Control and Reference Bias

Initial assessments showed variance in sequencing depth (Figure S6), a common precursor to allelic dropout. To ensure the reliability of our sister-species mapping, we performed a Reference bias check by quantifying the relationship between missing data and individual heterozygosity (H_O_). The lack of a significant negative correlation confirms that the use of the *O. eremita* genome did not artificially suppress diversity metrics in the more divergent *O. barnabita* samples. The reference-based pipeline recovered 6,150 loci and 27,364 SNPs, significantly outperforming the de novo approach, which yielded 4,405 loci and 9,053 SNPs (Supplementary Table 1). By utilizing a reference genome, we nearly tripled the SNP yield and increased locus recovery by approximately 33% while significantly reducing the proportion of missing data across the dataset. Principal Component Analysis (PCA) across both datasets revealed consistent broad-scale clustering, though the reference-based dataset offered tighter clusters and greater explanatory power due to the increased marker density and improved data completeness.

### 3.2. Genetic Diversity and Differentiation

Genetic diversity varied sharply across the study area. Populations in the Northeast (AF, BF) and the Southwest (SF) exhibited the lowest H_O_ and the highest positive F_IS_ values (0.604 to 0.785; Figure 2 and Figure S7, Table 2). The magnitude of subdivision was further highlighted by pairwise F_ST_ values. Differentiation ranged from a minimum of 0.021 (AF vs. BF) to a maximum of 0.360 (SF vs. BF). This extreme value for the SF population confirms its status as a highly isolated and divergent evolutionary unit.

**Fig. 2.**
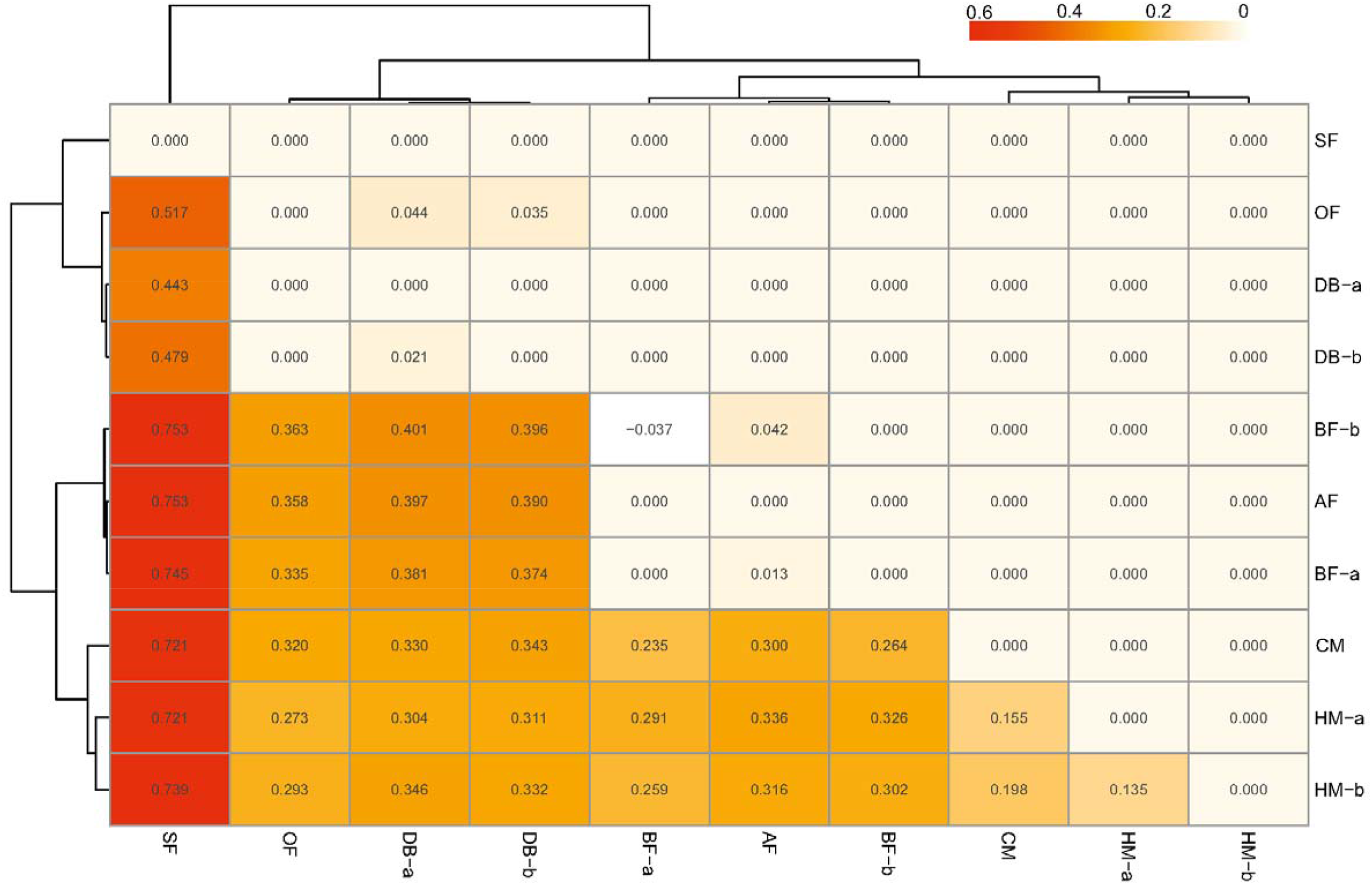
Pairwise population differentiation (F_ST_) among examined populations of Hermit beetles.

### 3.3. Population Structure and Phylogeny

The primary axis of variation (PC1) consistently isolated the SF population, a pattern that remained stable across the different datasets (Figure 3 and Figure S8,9). This genomic structure was further resolved by sNMF analysis which identified K=4 as the optimal number of ancestral lineages based on minimum cross-entropy (Figure 4). The Admixture bar plot for K=4 confirms four geographically distinct genetic lineages across Poland with the SF lineage showing minimal shared ancestry with neighbourhood populations (Figure 1 and Figure S8). Corroborating these findings, the neighbour joining tree revealed a remarkable long branch separating the SF clade from all other populations. (Figure 5). This phylogenetic distance coupled with the distinct clustering patterns observed across the multi - panel PCA (Figure S9) reinforces the hypothesis of a long-term isolation or a unique evolutionary trajectory for the Lower Silesian Forest population.

**Fig. 3.**
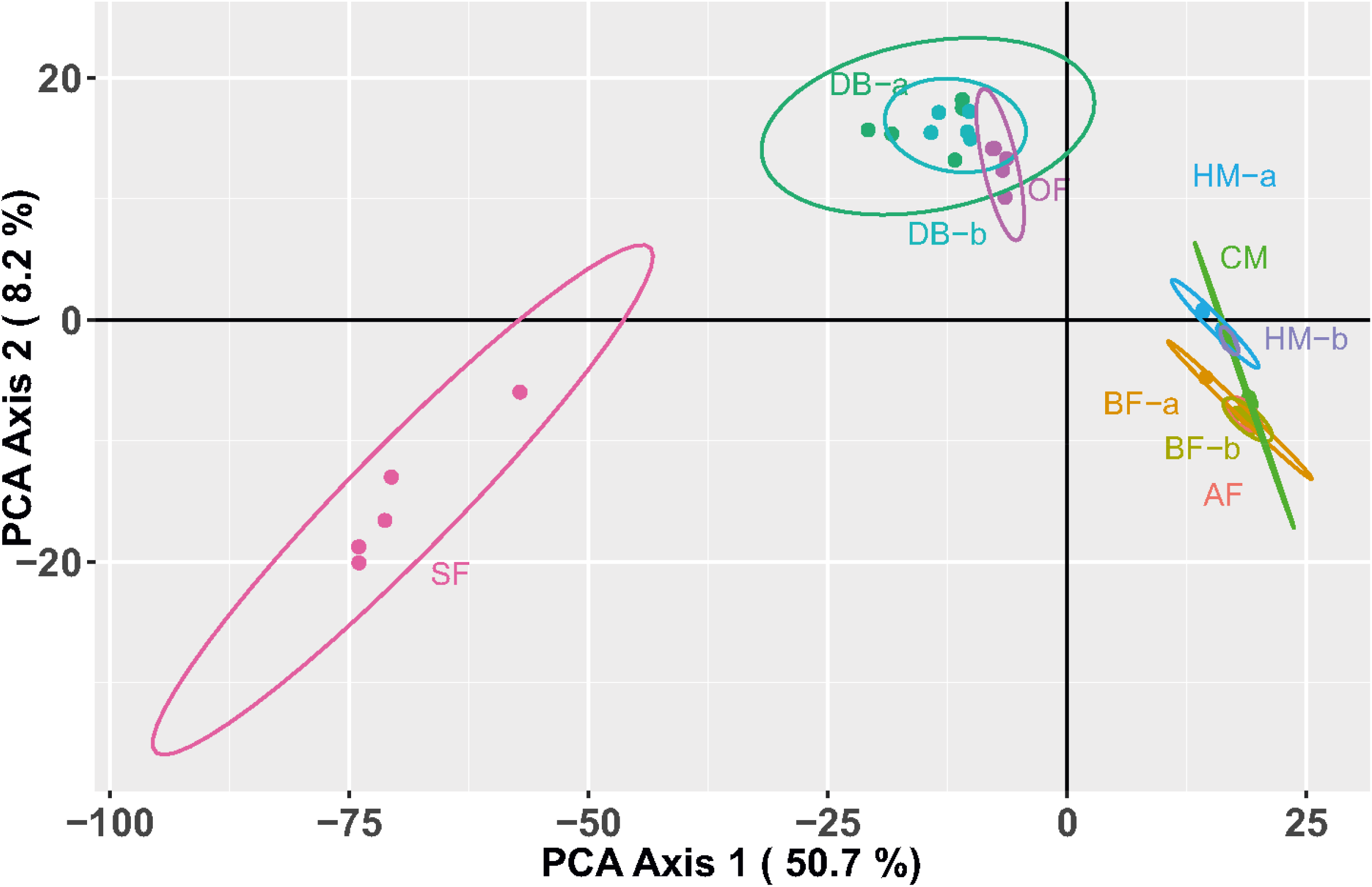
Principal Component Analysis (PCA) of 49 *O. barnabita* individuals (Reference-based dataset). Axes 1 and 2 depict the two dimensions explaining the most variance. Confidence ellipses (95%) are drawn around the population clusters.

**Fig. 4.**
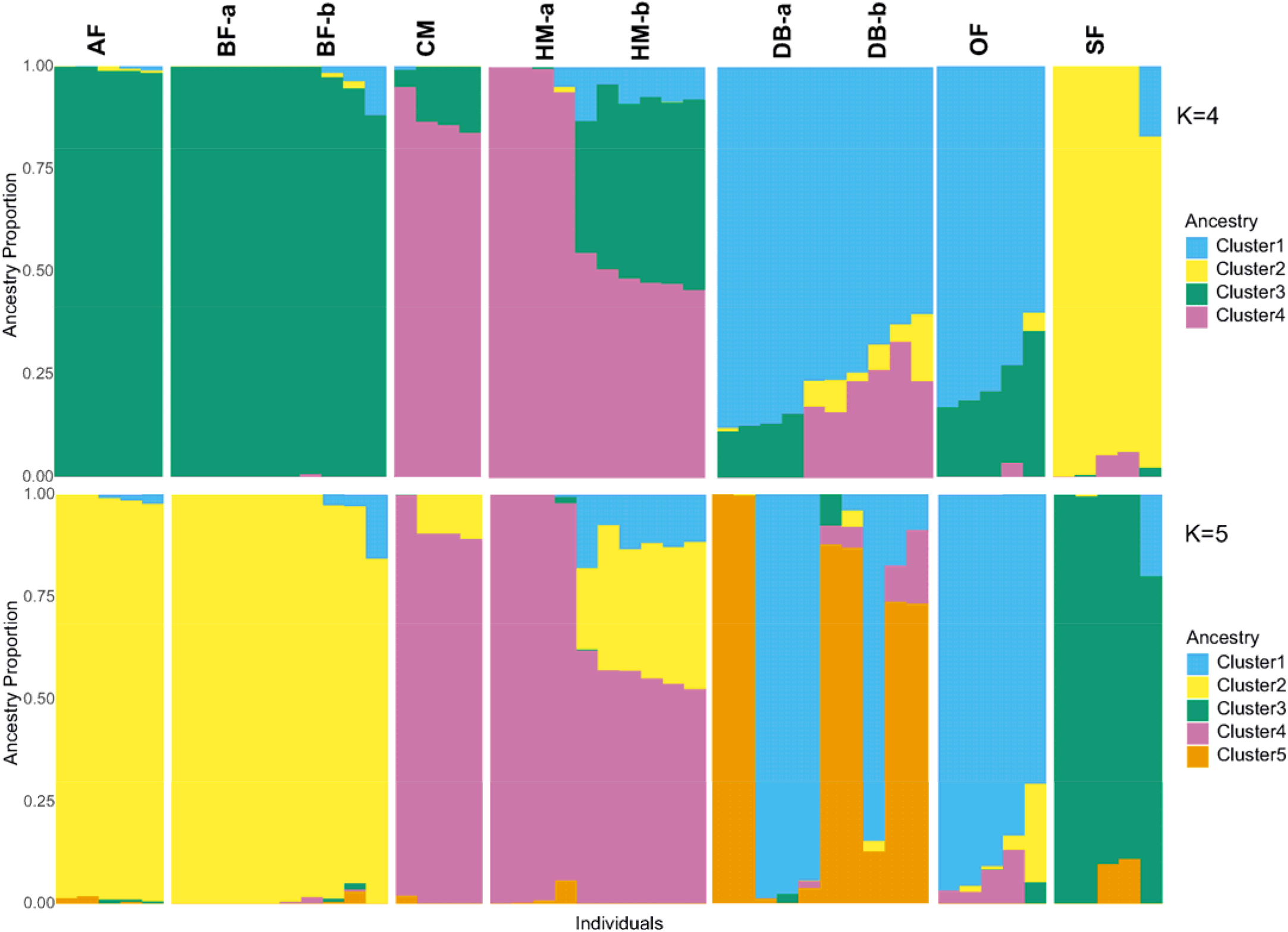
Admixture coefficients for K=4 ancestral clusters. Individuals are grouped and ordered by their designated population, showing the proportion of ancestry derived from each of the four clusters (represented by different colors). Below is the plot using K=5.

**Fig. 5.**
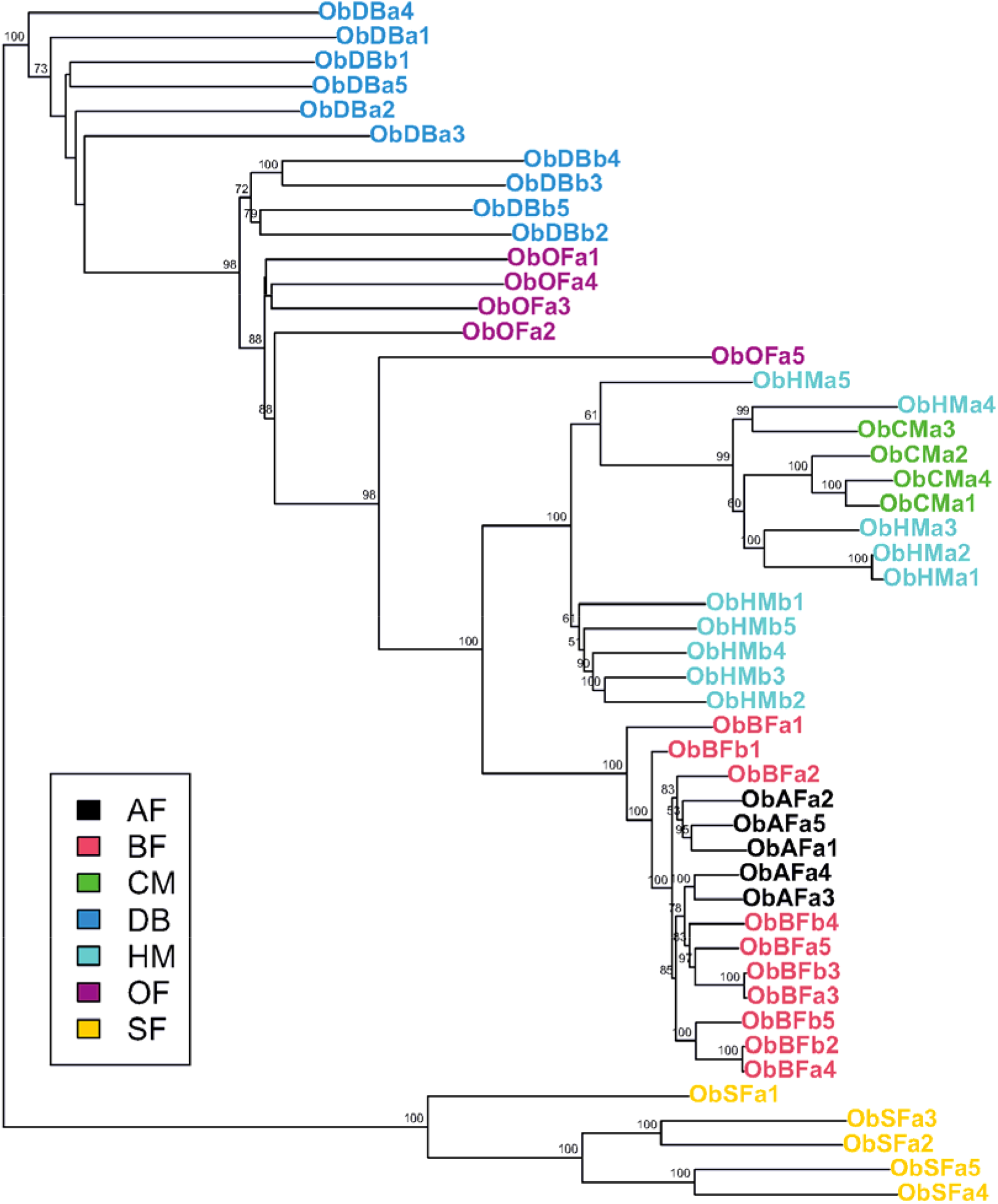
Neighbor-Joining (NJ) Tree of Osmoderma barnabita (using de novo dataset). This phylogenetic tree, constructed from a bitwise distance matrix, illustrates the relationships among the seven studied populations. The topology is characterized by a remarkably long branch that separates the SF clade from all other clusters, reinforcing the hypothesis that the Lower Silesian Forest population has a unique evolutionary trajectory or has undergone significant long-term isolation

### 3.4. Genomic Erosion and Adaptive Potential

We found a striking linear relationship between neutral genetic diversity and adaptive potential (R2=0.986, p<0.001; Figure S10). This near-perfect correlation indicates that the demographic processes, specifically genetic drift in small, fragmented forests are simultaneously depleting the standing genetic variation required for long-term persistence. Populations in the Northeast (AF, BF) showed the most severe signatures of genomic erosion, with H_e_ values dropping below 0.05 for both adaptive and neutral markers. In these sites, the "adaptive toolbox" has been severely affected by drift (Figure S10). In contrast, the southwestermost populations (DB, SF) maintained significantly higher diversity (H_e_ >0.20). In these populations, neutral diversity consistently exceeds adaptive diversity, a pattern expected under natural selection. In the eroded populations, however, the overwhelming force of drift has leveled the genomic landscape, removing the signal of selection entirely (Figure S10).

### 3.5. Landscape Patterns of Migration

EEMS analysis identified significant deviations from a simple isolation-by-distance model (Figure 6, Figure S11). A prominent genetic barrier (low effective migration) restricts gene flow through the central-western and eastern portions of the range. Higher historical connectivity was detected in the northeast and south-central regions (cyan/blue hues). The effective diversity surface (q-rates) revealed a strong longitudinal gradient. The western populations (SF, OF) act as high-diversity hotspots and stable genetic reservoirs, while the eastern populations function as diverse sinks with limited immigration.

**Fig. 6.**
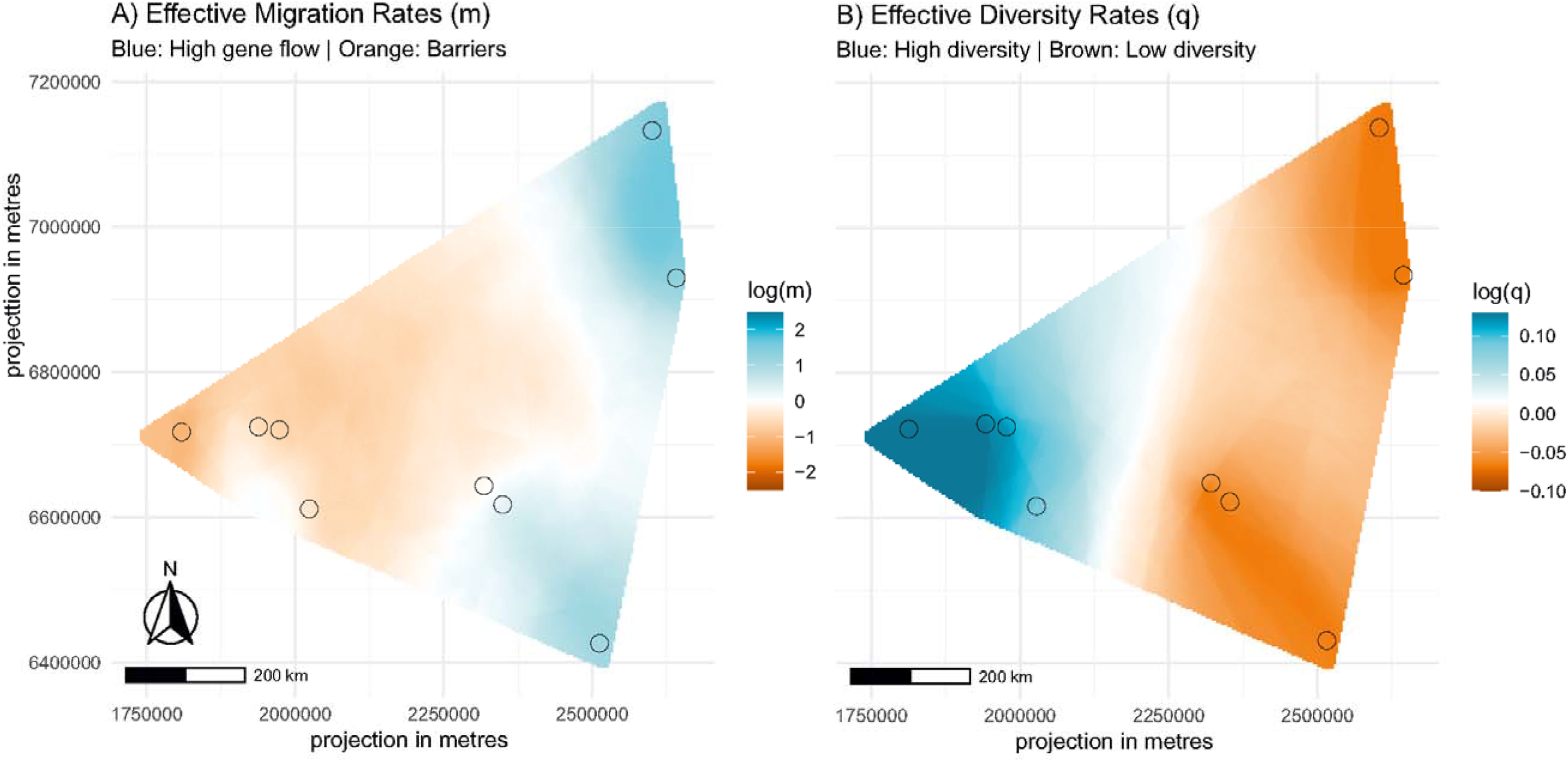
Estimated Effective Migration Surfaces (EEMS) for *Osmoderma barnabita*. (Reference based dataset) (A) Posterior mean effective migration rates (m) on a log10 scale. Cyan regions denote "corridors" of higher-than-average gene flow, while orange-to-brown contours identify spatial barriers where migration is significantly restricted relative to the global mean. (B) Posterior mean effective diversity rates (q) on a log10 scale. Cyan indicates areas of elevated genetic diversity (expected heterozygosity), while orange/brown identifies regions with lower-than-average diversity. All surfaces were interpolated using a 250-deme grid. Geographic coordinates are projected from WGS84.

### 3.6. COI Barcoding

BLAST analysis of the COI barcode sequences confirmed that all individuals, including those from the highly divergent SF population, belong to *Osmoderma barnabita* (sequence identity was up to 99.4% to sequences from Croatia (AJ880684) and Slovakia (AM423160), whereas the closest *O. eremita* from Germany (AM423156) had 91.0%).

## 4. Discussion

The primary discovery of this research is the profound genetic subdivision of *Osmoderma barnabita* populations in Poland, characterized by a massive genomic chasm between the westernmost population in the Lower Silesian Forest and the remaining populations, especially belonging to eastern and central clusters. Our integrated approach consistently identified four major genetic lineages and that there is limited or no gene flow amongst these. These findings mandate a reevaluation of current conservation frameworks (Oleksa et al., 2012); the Polish metapopulation should be delineated into at least three Evolutionary Significant Units (ESUs) and managed as independent Management Units (MUs), despite a lack of mitochondrial differentiation. The congruence between our *de novo* and reference-based pipelines confirms the reliability of these findings. While the *O. eremita* reference genome helped capture the deep divergence of the western relict, the *de novo* results proved that the observed subdivision is a biological reality, not a bioinformatic artifact. It is also an important finding, as so far no previous studies have proved the reliability of various bioinformatic approaches in determining genotypes of saproxylic beetles, as it was done for other insects (e.g. Deng et al., 2022). It is an important finding, as methods such as ddRADseq or Genotyping-by-sequencing are cost-effective and require a lower amount of DNA, than whole genome sequencing, and therefore could be easily adapted for e.g. population and ecological studies or monitoring of Hermit beetles, and other saproxylic beetles.

### 4.1. The Genomic Advantage in Saproxylic Conservation

Previous studies using mitochondrial DNA or microsatellites provided valuable snapshots of the Hermit Beetle’s phylogeographic history and small-scale isolation (Audisio et al., 2009; Oleksa et al., 2015, Landvik et al., 2015, Melosik et al., 2020). These studies proved phylogenetic distinctiveness of Hermit beetles in Europe (Audisio et al., 2009), low genetic polymorphism of northeastermost populations (Landvik et al., 2015), and highly limited dispersal in local scale (Oleksa et al., 2015, Melosik et al., 2020). However, these markers lack the resolution to detect fine-scale population structure or adaptive potential. Our use of ddRADseq represents a shift toward genomic standards in saproxylic beetle research (Kajtoch et al., 2022). This high-resolution approach clarifies deep subdivision and highlights the limitations of using single-locus mtDNA to define conservation units in species with complex post-glacial histories (Taberlet et al., 1998, Krovi et al., 2025).

### 4.2. Biogeographic drivers of population structure

#### 4.2.1. The Isolation of the Lower Silesian Forest population

The Lower Silesian Forest population of *O. barnabita* represents the most significant genetic discontinuity in the study, isolated on the primary PCA axis with extreme pairwise F_ST_ values (0.250-0.360). This divergence suggests a different refugial origin compared to the rest of the Polish populations which was proposed in other high-fidelity saproxylic specialists like *Lucanus cervus and Rosalia alpina* (Cox et al., 2019; Drag et al., 2015, 2018; Krovi et al., 2025). While its western position might suggest an affinity with *O. eremita*, the presence of *O. barnabita* mitochondrial haplotypes points toward a more complex history, potentially a hybrid origin with an introgressed genome. Alternatively, the SF population may have undergone long-term functional isolation. Although oak-lime woodlands are widespread in Silesia and lack physical barriers to migration (Kadej et al., 2014), the continuity of hollow tree habitats has been severely reduced. This creates a functional isolation as well. The exceptionally high inbreeding coefficient (F_is_ = 0.604) in the SF population is consistent with the sedentary life-history of *Osmoderma*, where dispersal is often limited to less than 200 m from the natal tree (Ranius & Hedin, 2001). Furthermore, the persistence of *O. barnabita* mitochondrial haplotypes alongside a highly divergent nuclear genome suggests a complex history of secondary contact or incomplete lineage sorting. Rather than representing a degraded unit, the SF population acts as a relictual lineage preserving unique alleles that are absent in the eastern clusters, making its preservation a mandate for maintaining the species’ overall adaptive breadth.

#### 4.2.2. Connectivity in the Eastern and Central Lineages

The populations follow a structure shaped by historical continuity and recent breakdown: The Białowieża (BF) and Augustów (AF) forests share a single ancestral cluster with minimal differentiation (F_ST_≈0.021). This confirms the Podlasie region as a primary stronghold for the Eastern European *O. barnabita* clade, where large, contiguous forest tracts still facilitate the gene flow necessary to buffer against genetic drift (Oleksa et al., 2013). This interpretation is a bit uncertain due to the observed low heterozygosity of BF and AF populations. It could be a result of the phylogeographic history of the Hermit beetle expansion, which reached northern areas later than e.g. forests in Silesia or the Carpathians, spreading most likely from the Balkans (Audisio et al., 2009). Also, beetles from Baltic states and Finland have reduced mitochondrial diversity (Landvik et al., 2015). It could also be connected with climate changes, which promote warm-adapted oaks and limes, that replace cold-adapted trees like spruces in well known primeval Białowieża Forest (Jaroszewicz et al., 2019). These two are not mutually exclusive, and both might lead to founder effect reducing the genetic diversity of Hermit beetles in their northeastermost range.

A second major lineage occupies the Carpathian (CM) and Holy Cross (HM) Mountains. The detected admixture in the HM population points to a historical forest bridge along the eastern forest belt, a known migration corridor for various saproxylic taxa moving between the Carpathians and the Central Polish uplands (Kajtoch et al., 2016). However, while habitat networks and wildlife corridors are increasingly promoted as a priority conservation strategy (MacDonald, 2003; DeFries et al., 2023), their effectiveness requires empirical evaluation in most systems. Our genomic results showed that the rising F_is_ values and the clear lineage separation suggest that while these corridors were historically functional, they are currently failing to maintain connectivity.

The Oder Valley (OF) population represents a third lineage. Interestingly, the nearby Barycz Valley (DB) shows admixture from both the OF and the divergent SF clusters, suggesting that while some gene flow persists in Silesia, it is insufficient to bridge the "chasm" created by the isolation of the Lower Silesian Forests. Silesia, with adjacent Czech and German forests, is a well-known zone of admixture for saproxylic beetles, e.g. for recently examined red flat bark beetles *Cucujus cinnaberinus* (Sikora et al., 2023). This part of Europe likely acts as a transient zone for the west and east clades of forest species, including deadwood-dwelling Coleoptera (Krovi et al., 2025), therefore is important for studying evolutionary processes and ecological adaptations of beetles used to different climatic and environmental conditions. This zone is also on the borderline between intensively transformed and managed forests in the west and more natural forests in the east, which leave signs in genetic signatures of saproxylic beetles (Amer et al., 2025).

### 4.3. Conservation and Management Implications

Our high-resolution map reveals a species caught between ancient biogeographic history and modern human impact. The extreme differentiation of the SF unit, coupled with the poor dispersal capacity of Hermit Beetles (Oleksa et al., 2015, Melosik et al., 2020), makes it a priority for isolated management. These results have immediate practical applications for re-introductions and translocations. To avoid outbreeding depression or localized adaptive failure, source populations must be matched to their respective evolutionary lineages. For example, moving individuals from the semiboreal northeastern lowlands to the humid, montane beech forests of the south could lead to a mismatch in environmental fitness. This attention to detail in lineage matching is vital because the conservation of *O. barnabita* is inherently linked to the maintenance of a broader forest biodiversity. It stems from the species’ stringent microhabitat requirements as recent evidence suggests that even within suitable climates, the beetle is limited by a lack of veteran trees with specific wood-mould characteristics and stable microclimates, which are increasingly rare in managed forest landscapes (Smolis et al., 2023). Recent evidence highlights the species as a critical umbrella or a surrogate taxon (Smolis et al., 2026). Because the presence of a beetle is a significant predictor of the tree-related microhabitats, the loss of a genetic lineage could signal the degradation of the complex micro habitat communities they indicate. Therefore, a lineage-based strategy, similar to that used for the Great Capricorn Beetle (*Cerambyx cerdo*), is essential (Drag & Cizek 2015) to preserve the genetic integrity. Protecting the genomic integrity of these beetles requires not only preserving individual trees or larger aggregations of them but also maintaining the "milestones" between local populations necessary to prevent the extinction of these evolutionary lineages.

## 5. Conclusion

This population genomic assessment reveals a striking dichotomy: a historical pattern of post-glacial colonization that forged four distinct genetic lineages, now overlaid by the severe signatures of modern habitat fragmentation. The west-to-east genetic differentiation observed in Polish *Osmoderma barnabita* aligns with the broader phylogeographic paradigm for European saproxylic beetles (Kajtoch et al., 2022; Krovi et al., 2025). This study is focused on Polish populations, although it is likely that distinct lineages also exist in surrounding forests in Germany, the Czech Republic or Austria. The extension of the genetic study to other populations is required and should be linked with conservation actions on this charismatic species (e.g. LIFE Osmoderma https://www.osmoderma.lt/). The Lower Silesia population represents a critical evolutionary legacy that likely reflects a unique expansion episode or a complex hybrid origin. Given its extreme isolation and high inbreeding, the conservation of *O. barnabita* must immediately prioritize this Evolutionary Significant Unit (ESU) to prevent the loss of its unique genetic heritage. Furthermore, our data demonstrate that conserving *O. barnabita* requires maintaining not just individual trees but the entire ecosystem with hollow tree habitats that preserve connectivity among subpopulations in regions occupied by beetles. The open question is whether conservation of Hermit beetles should also restore connectivity between deeply divergent evolutionary lineages, as it could lead to loss of their adaptive potential and cause outbreeding depression. Considering admixed populations (e.g. Holy Cross Mountains), decisions on which populations could be connected or should not be, need to be preceded by genetic evaluation. This is especially important in the case of translocation, reintroduction, or restoration of natural wooded bridges among distinct populations. Without "stepping-stone" corridors of veteran trees to restore gene flow inside regional populations (e.g. within Silesia and Podlasie regions), the adaptive potential of these isolated forest relicts will continue to erode, leaving them vulnerable in an increasingly fragmented temperate ecosystem.

## Ethical statement

Collection of protected beetle species and in protected areas was executed according to permissions granted by Polish Ministry of Environment (DOP-1.61.61.2021.TP 1516071.4994962.3977511 and DOP-1.61.61.2021.TP 1516071.4994586.3977515), General Directorate of Environmental Protection (DZP-WF.6401.68.2021.AS) and Regional Directorates of Environmental Protection in Białystok (WPN.6205.74.2021.MM), Rzeszów (WPN.6205.116.2021.MP.2 and WPN.6205.114.2021.ŁL.2), Kielce (WPN.I.6205.1.49.2021.EJP.2) and Wrocław (WPN.6205.143.2021.AR).

## Data Availability Statement

The genomic data generated in this study are available across public repositories. The raw de-multiplexed ddRADseq reads are deposited in the NCBI Sequence Read Archive (SRA) under BioProject PRJNA1438437 (Submission ID: SUB16065778). Individual mitochondrial nucleotide sequences are available in GenBank under accession numbers PZ336326–PZ336344. Additionally, the filtered SNP datasets (VCF format) used for population genomic analyses are archived in the RepoOD repository (https://repod.icm.edu.pl/dataverse/isez).

## Acknowledgements and Funding

This research was funded by the National Science Centre, Poland (NCN grant UMO-2021/43/B/NZ9/00991). The authors are sincerely grateful to the Polish nature-protection authorities for granting permission to conduct field work in protected forest complexes and to collect protected species, and to the Polish State Forests National Forest Holding, particularly the Regional Directorates of State Forests in Krosno, Radom, Wrocław, and Białystok, as well as the local foresters.

## Supplementary files

**Figure S1.**
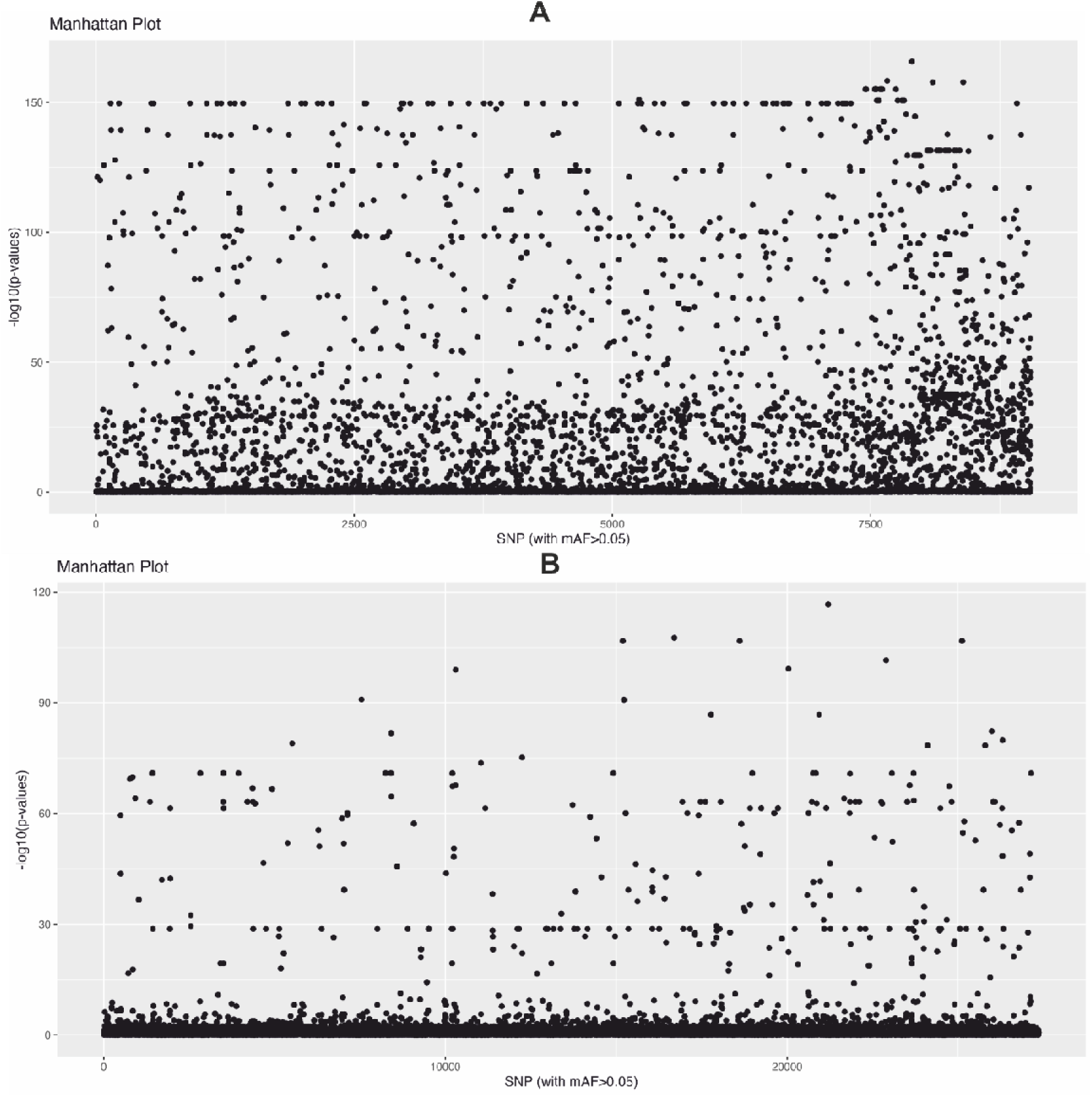
Genome-wide scan for signatures of selection/outlier detection in Osmoderma barnabita for De Novo Pipeline (A) and Reference based Pipeline (B). The Manhattan plot illustrates the distribution of p-values for 27,364 SNPs across the genomic scaffolds of the O. eremita reference. Each point represents an individual locus. The X-axis represents the physical position of SNPs along the reference scaffolds (organized by scaffold ID). The Y-axis displays the negative decadic logarithm of the p-values (−log10(p)), where higher values indicate a greater departure from neutral expectations. The horizontal dashed line represents the significance threshold (α=0.05) after applying a Benjamini-Hochberg (BH) correction for multiple testing. Points exceeding this threshold (2,920 loci) are considered putative outliers under directional selection and were sequestered from the neutral dataset to avoid confounding demographic inferences.

**Figure S2.**
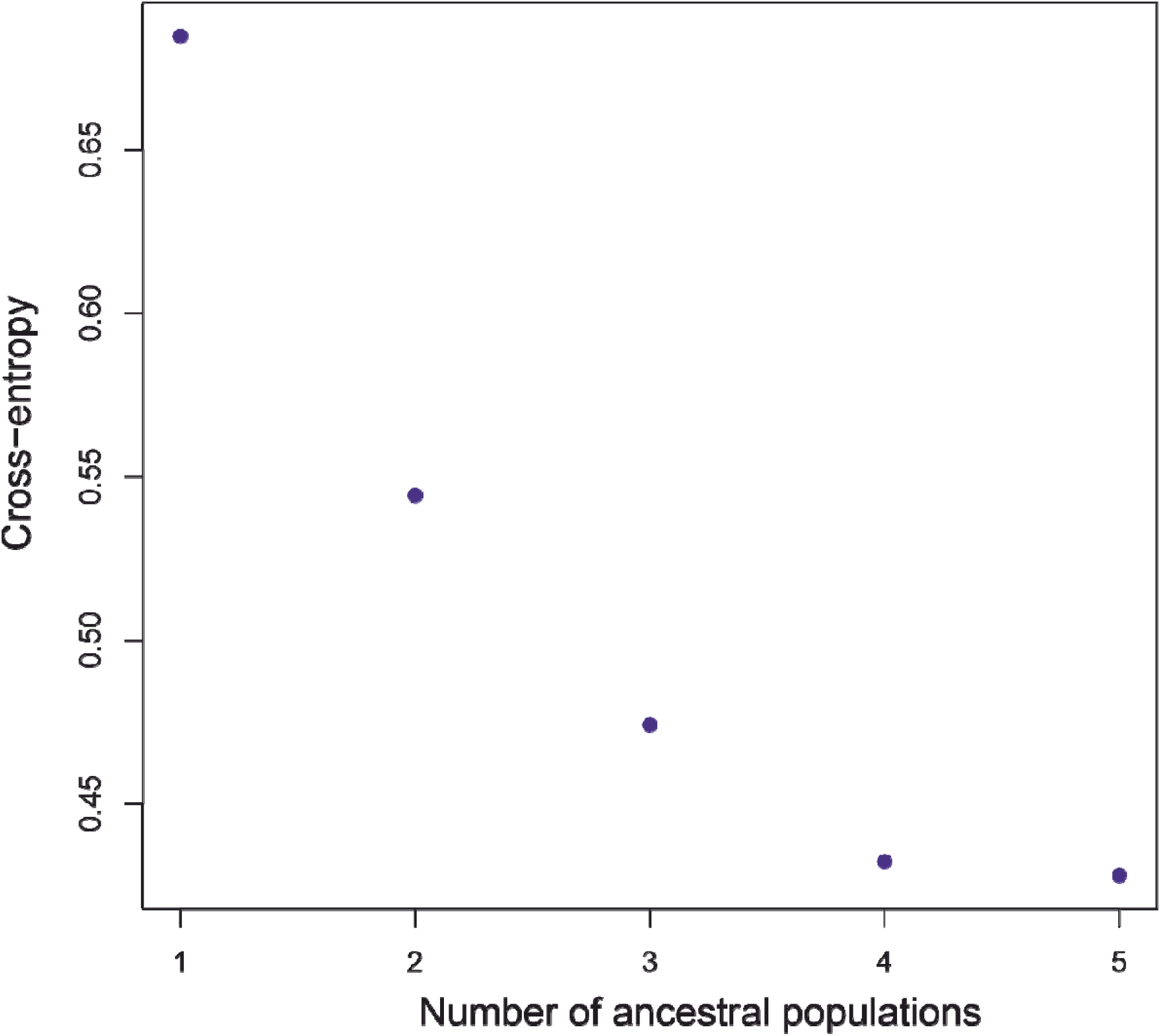
Plot of cross-entropy values calculated by sNMF for K=1 through K=7. Reference basedpipeline.

**Figure. S3.**
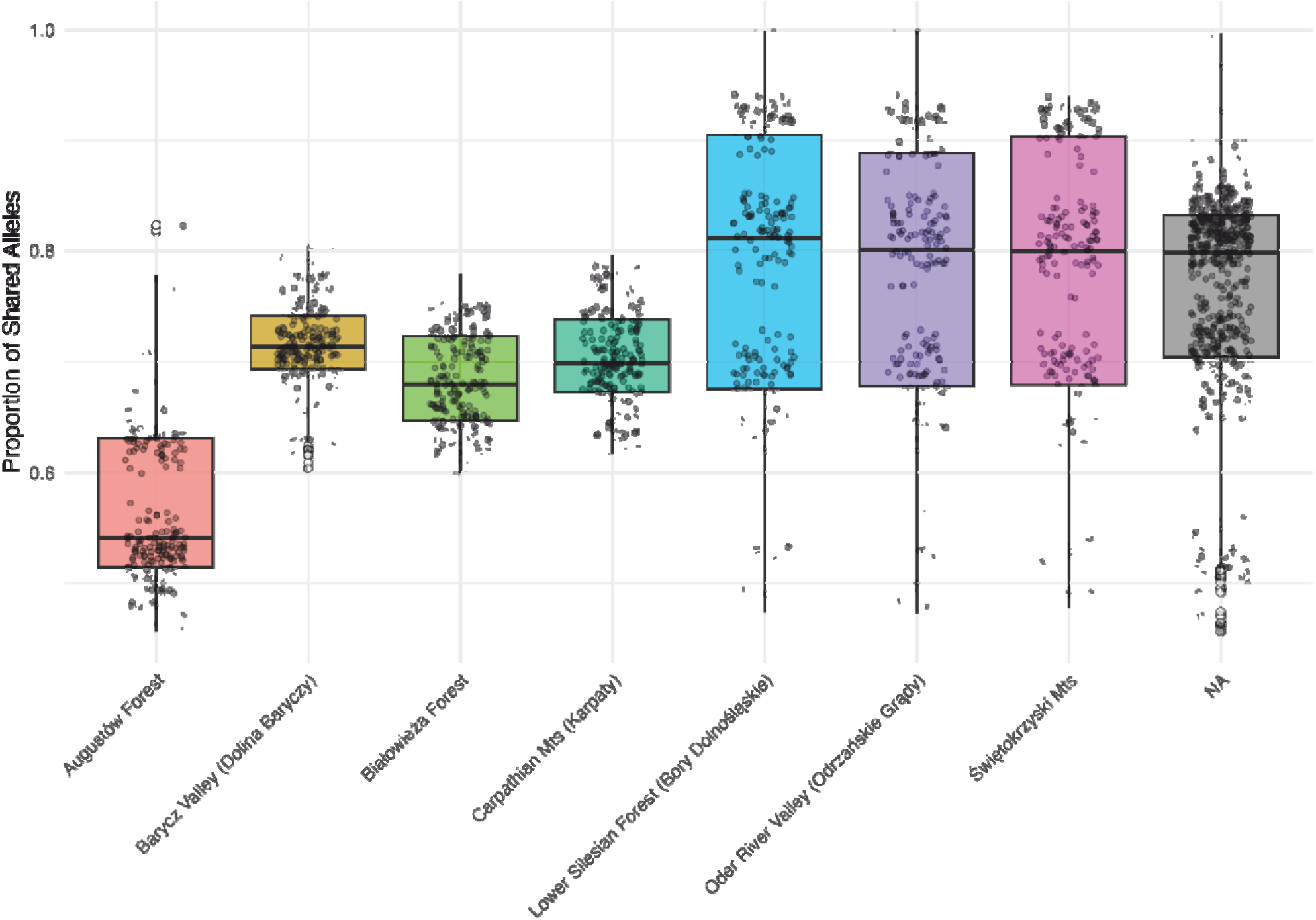
Population genetic relatedness in Osmoderma barnabita. Boxplots represent the distribution of the proportion of shared alleles between pairs of individuals among all forests. Gray jittered points represent individual pairwise comparisons.

**Figure S4.**
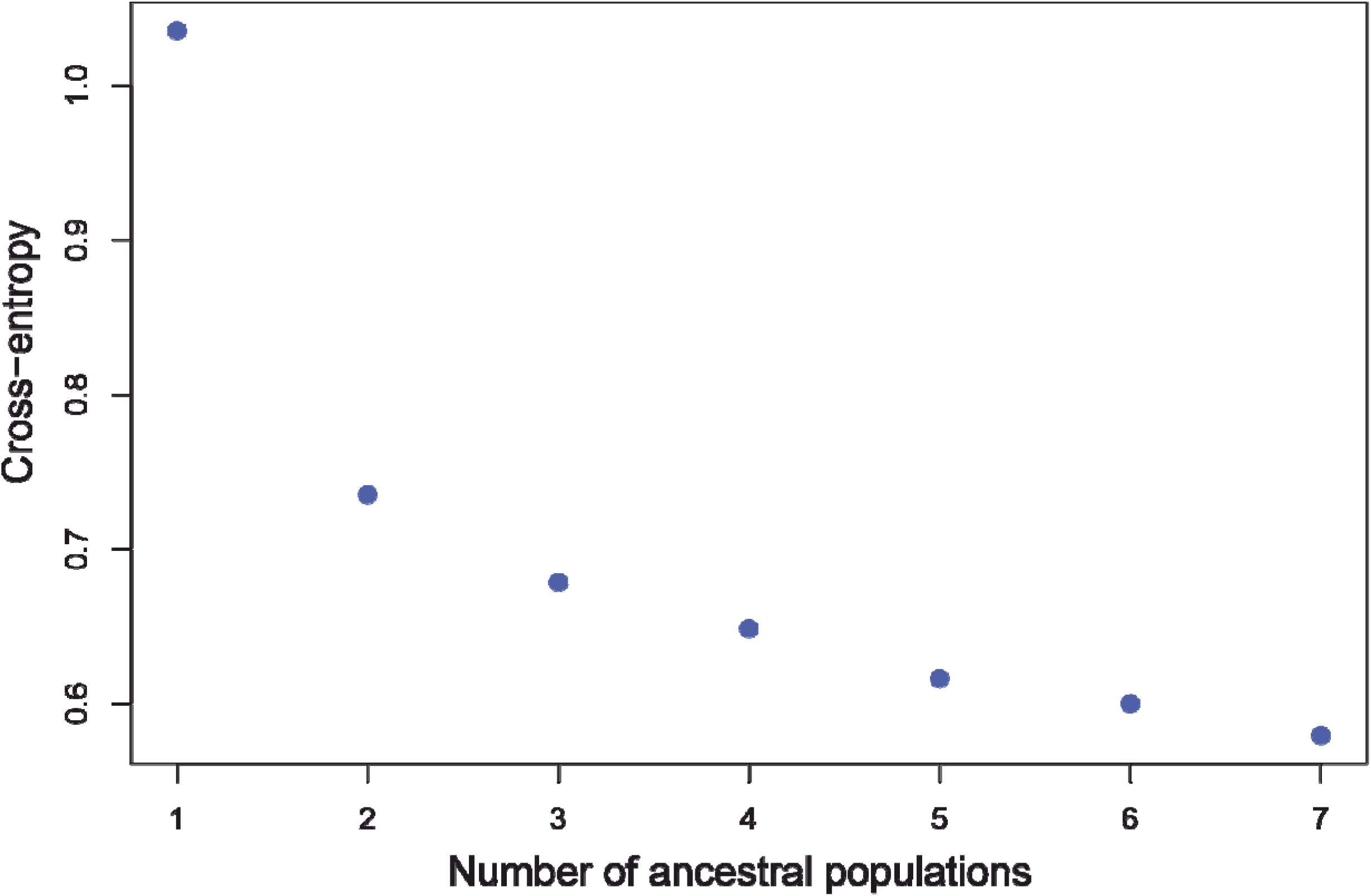
Plot of cross-entropy values calculated by sNMF for K=1 through K=7. De novo based pipeline.

**Figure S5.**
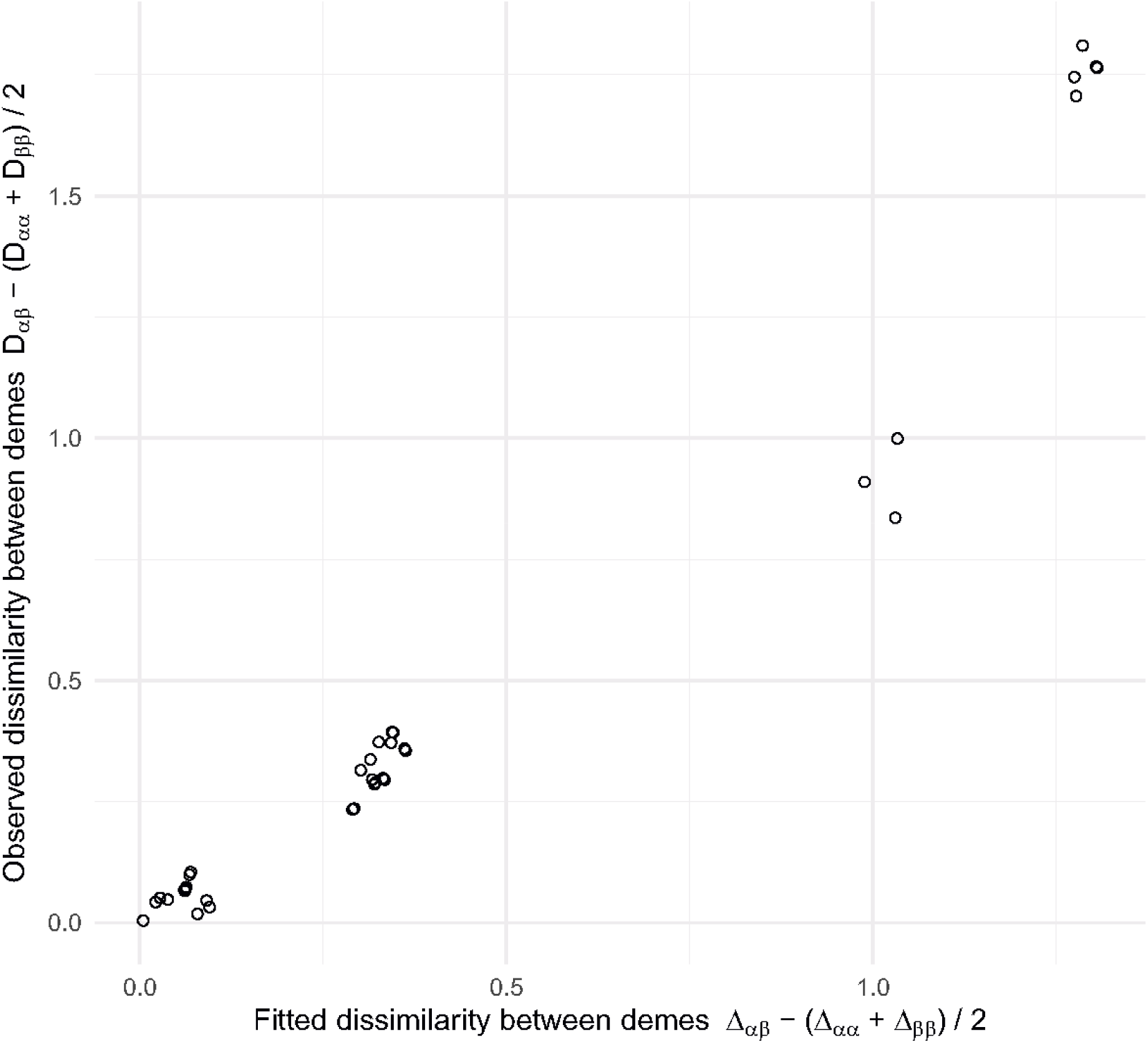
Model Goodness-of-Fit. Comparison of observed vs. fitted genetic dissimilarities between pairs of sampled demes. Each point represents a pair of demes (alpha, beta). The x-axis shows the dissimilarity (Delta) predicted by the EEMS model, while the y-axis shows the observed genetic dissimilarity (D). The tight alignment of points along the diagonal indicates that the model accurately captures the spatial genetic structure of the dataset. Singleton demes are excluded from the visualization to ensure accurate within-deme variance calculations

**Figure S6.**
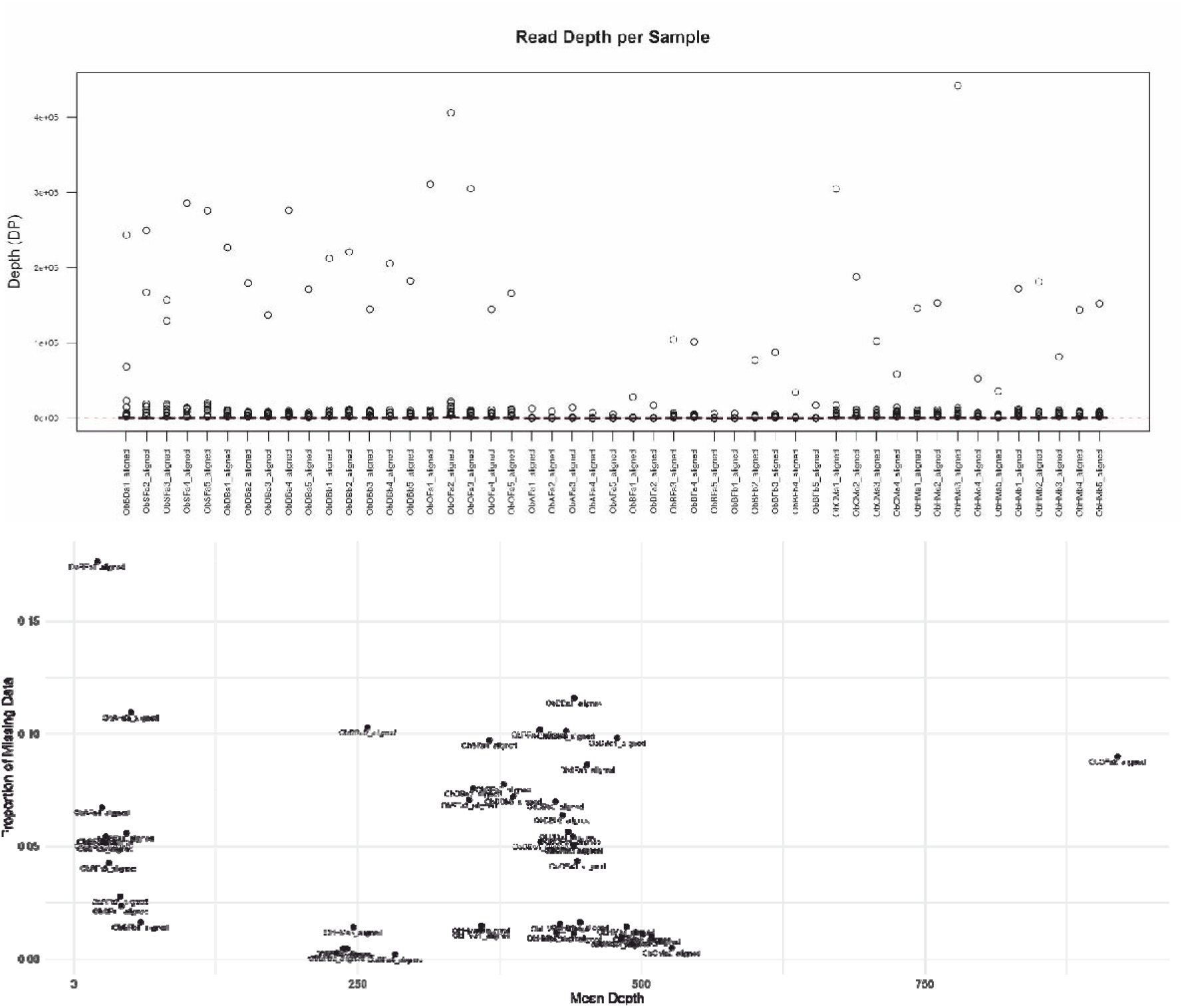
A - Sequencing Depth Distribution. This figure shows the read depth (DP) (on the y-axis) across all sampled individuals (x-axis). While median depth remains consistent across all the samples. B - Correlation between Missingness and Read Depth. This plot evaluates sample quality. Samples clustered at the bottom right indicate high-confidence sequencing.

**Figure S7.**
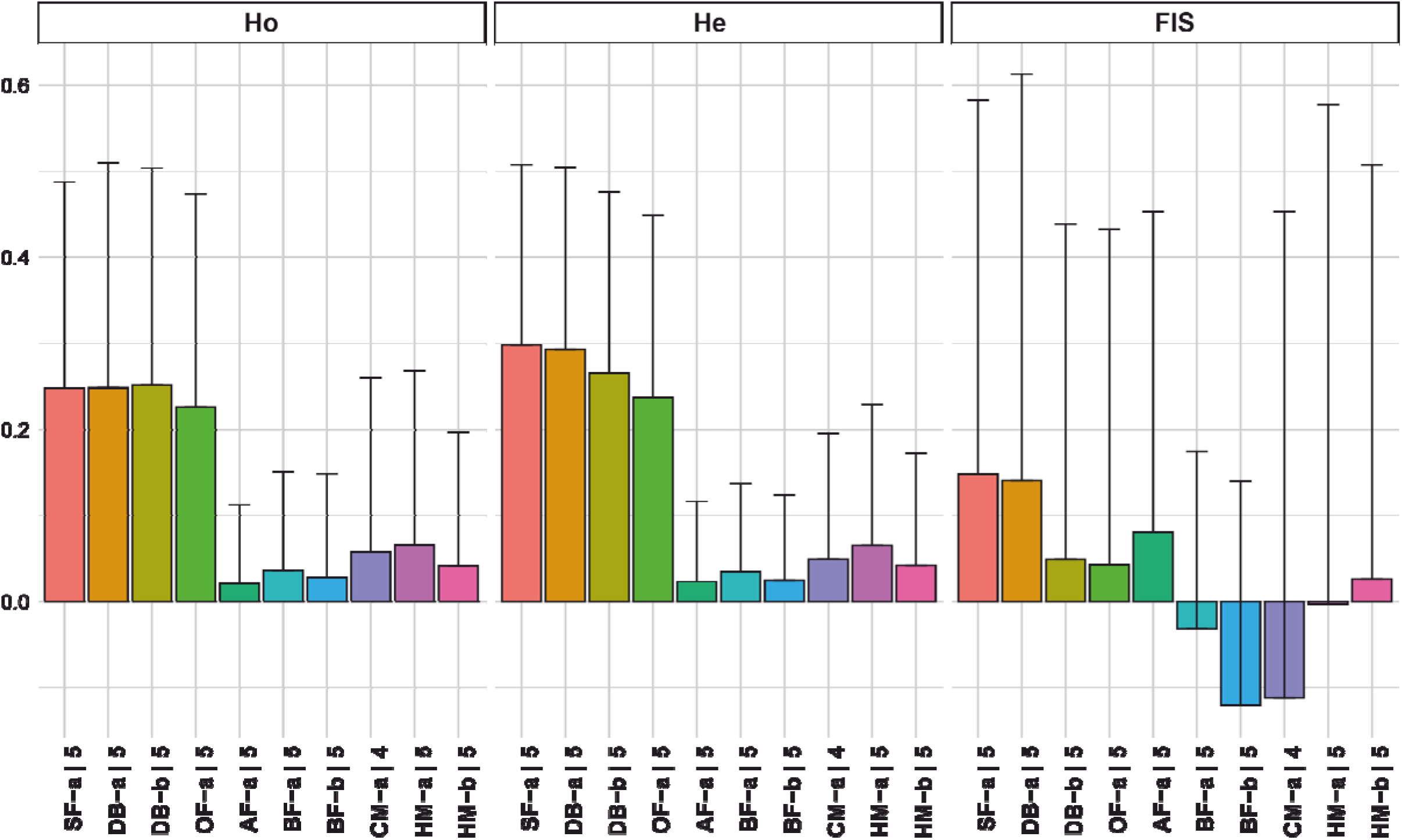
Population Genetic Diversity and Inbreeding Coefficients. Plot shows the genetic diversity metrics across ten sampled populations (labeled on the x-axis with associated sample sizes). The data is categorized into three panels: Observed Heterozygosity (Ho), Expected Heterozygosity (He), and the Inbreeding Coefficient (FIS). Each bar represents the mean value for a specific population, with vertical error bars indicating one standard deviation from the mean.

**Figure S8.**
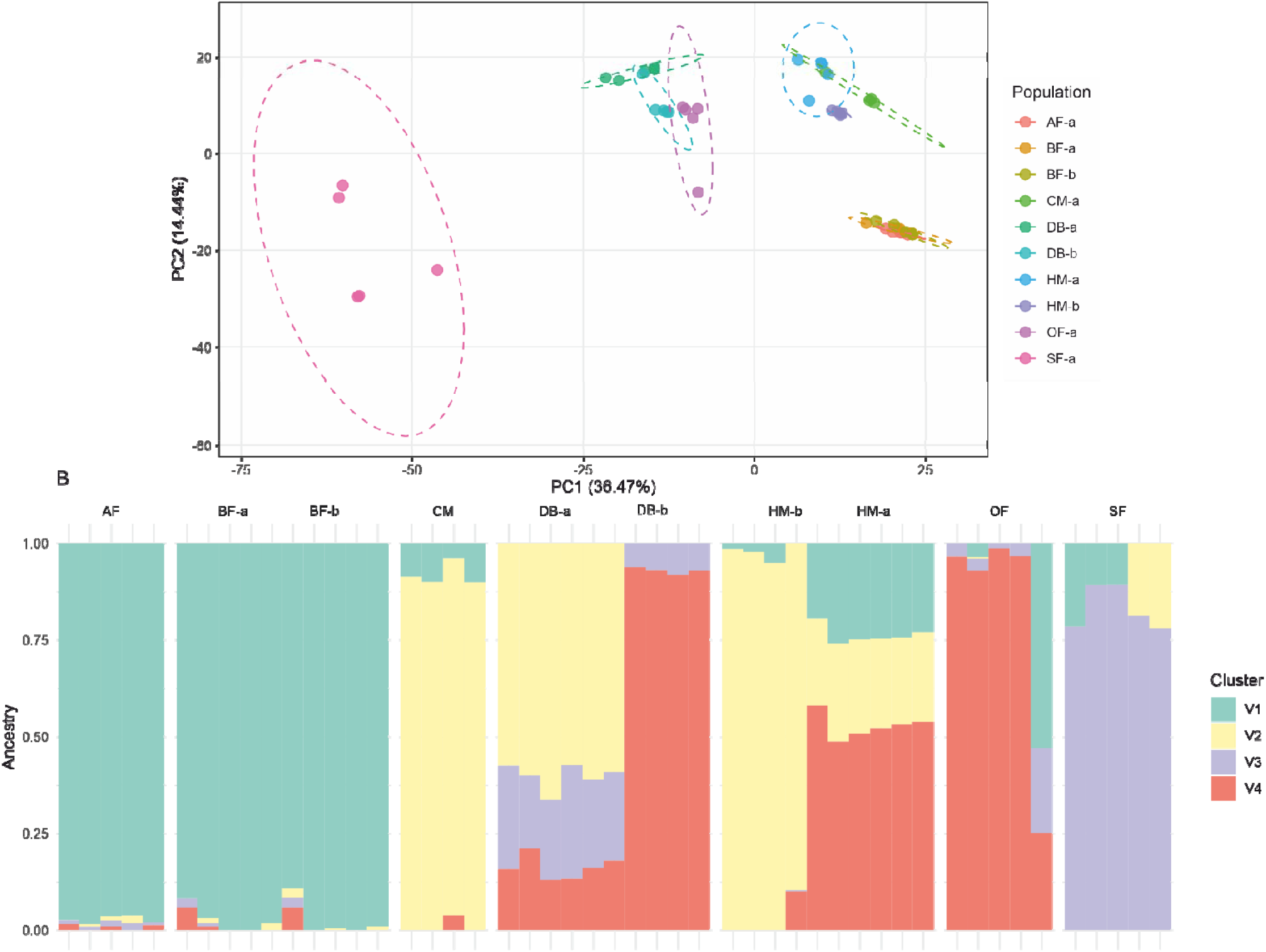
Population structure of Osmoderma barnabita based on de novo pipeline A - PCA, B - Admixture.

**Figure S9.**
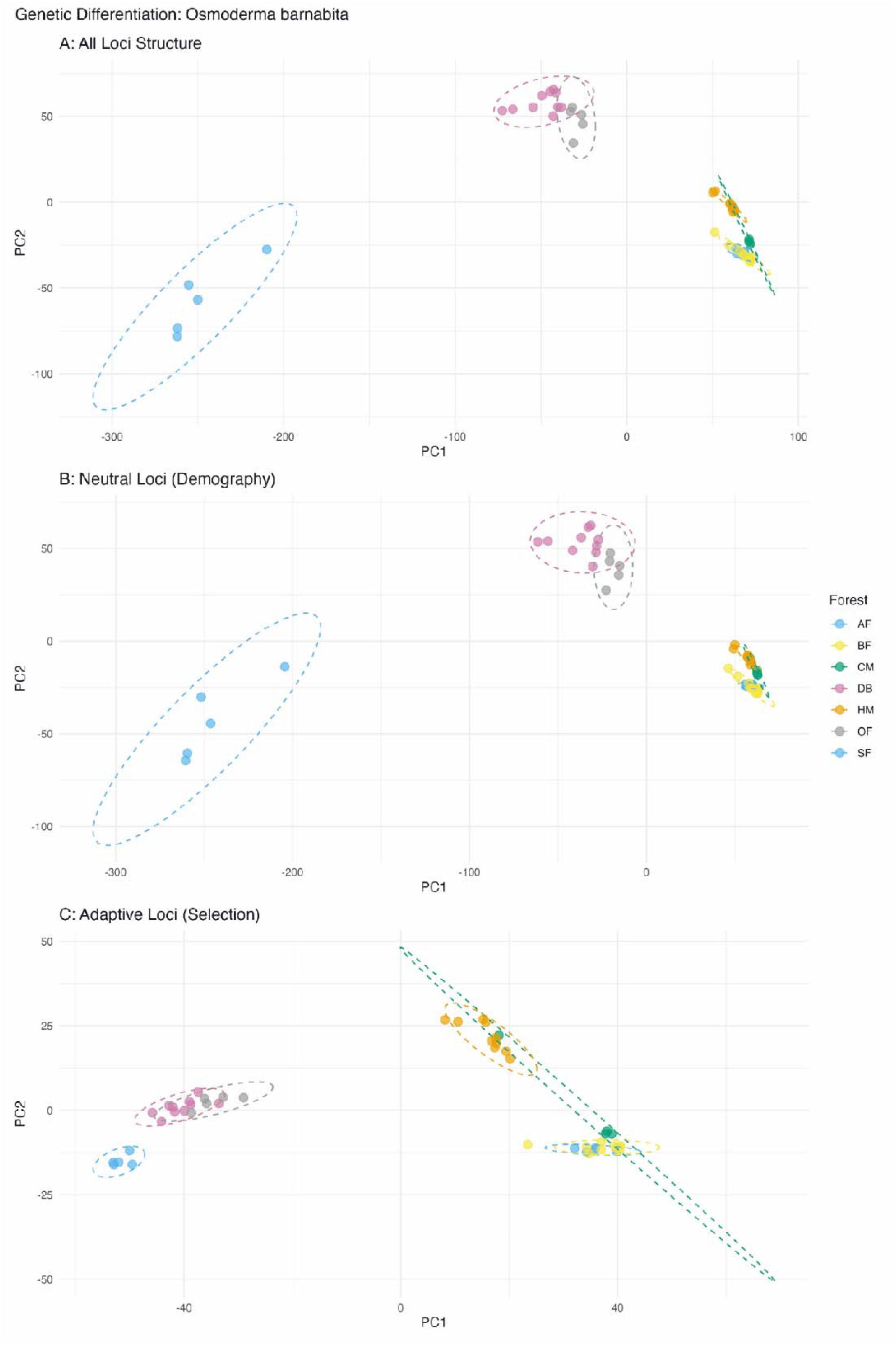
The genetic structure of Osmoderma barnabita across various forest populations using Principal Component Analysis. Across all panels, dashed ellipses represent 95% confidence intervals for each population cluster.

**Figure S10.**
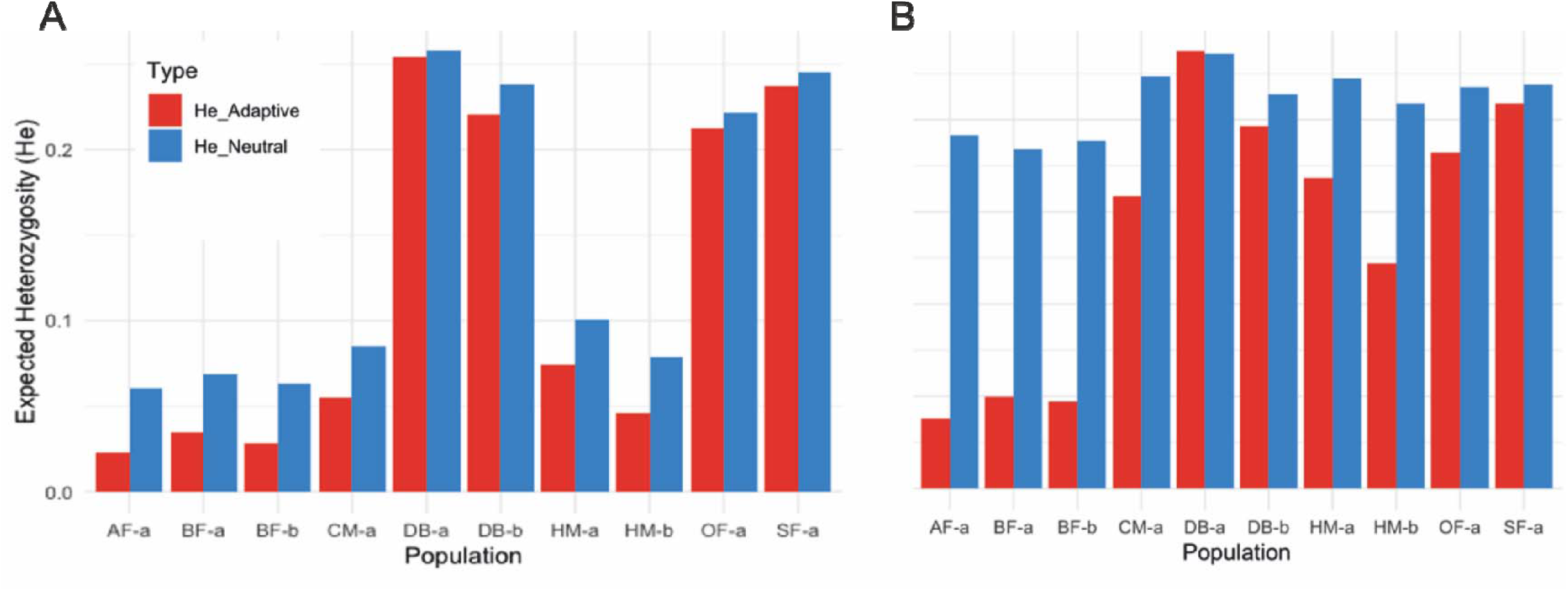
Plots visualizing neutral and adaptive heterozygosity distribution among examined populations of Osmoderma barnabita based on Reference Based Pipeline (A) and de novo (B).

**Figure S11.**
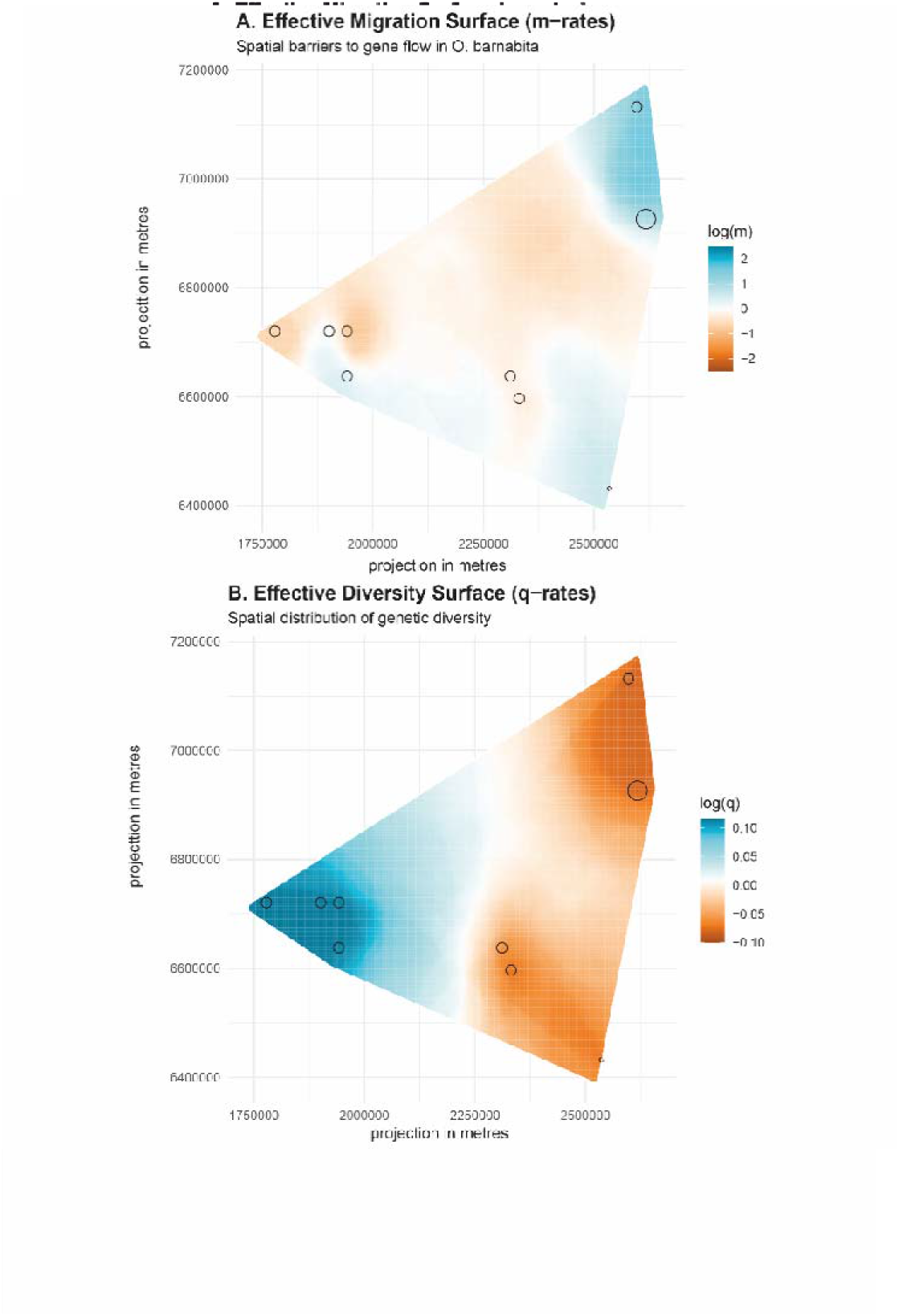
Estimated Effective Migration Surfaces (EEMS) for Osmoderma barnabita. (Denovo based dataset) (A) Posterior means effective migration rates (m) on a log10 scale. Blue colors indicate regions of high gene flow (migration "highways"), while orange/brown colors highlight barriers where migration is restricted. (B) Posterior mean effective diversity rates (q) on a log10 scale. Blue indicates regions of higher-than-average genetic diversity; brown indicates lower-than-average diversity. Analysis was performed using 250 demes.

**Table S1.** Comparative summary of ddRADseq data processed using two pipelines (referenced based and de novo).

| reference based |  |  |  |  |  |  |  |  |  |
| --- | --- | --- | --- | --- | --- | --- | --- | --- | --- |
| sample | n_loci | present_loci | missing_loci | frequency_missing | sample | n_sites | present_sites | missing_sites | frequency_missing |
| ObSFa1_aligned | 6150 | 6147 | 3 | 0.0005 | ObBDa1_aligned | 27364 | 25816 | 1548 | 0.0566 |
| ObSFa2_aligned | 6150 | 6140 | 10 | 0.0016 | ObSFa2_aligned | 27364 | 25242 | 2122 | 0.0775 |
| ObSFa3_aligned | 6150 | 6044 | 106 | 0.0172 | ObSFa3_aligned | 27364 | 25435 | 1929 | 0.0705 |
| ObSFa4_aligned | 6150 | 6131 | 19 | 0.0031 | ObSFa4_aligned | 27364 | 24706 | 2658 | 0.0971 |
| ObSFa5_aligned | 6150 | 6142 | 8 | 0.0013 | ObSFa5_aligned | 27364 | 25006 | 2358 | 0.0862 |
| ObDBa1_aligned | 6150 | 6134 | 16 | 0.0026 | ObDBa1_aligned | 27364 | 24192 | 3172 | 0.1159 |
| ObDBa2_aligned | 6150 | 6127 | 23 | 0.0037 | ObDBa2_aligned | 27364 | 24580 | 2784 | 0.1017 |
| ObDBa3_aligned | 6150 | 6145 | 5 | 0.0008 | ObDBa3_aligned | 27364 | 25295 | 2069 | 0.0756 |
| ObDBa4_aligned | 6150 | 6149 | 1 | 0.0002 | ObDBa4_aligned | 27364 | 24678 | 2686 | 0.0982 |
| ObDBa5_aligned | 6150 | 6150 | 0 | 0 | ObDBa5_aligned | 27364 | 24549 | 2815 | 0.1029 |
| ObDBb1_aligned | 6150 | 6144 | 6 | 0.001 | ObDBb1_aligned | 27364 | 24595 | 2769 | 0.1012 |
| ObDBb2_aligned | 6150 | 6131 | 19 | 0.0031 | ObDBb2_aligned | 27364 | 25615 | 1749 | 0.0639 |
| ObDBb3_aligned | 6150 | 6150 | 0 | 0 | ObDBb3_aligned | 27364 | 25451 | 1913 | 0.0699 |
| ObDBb4_aligned | 6150 | 6147 | 3 | 0.0005 | ObDBb4_aligned | 27364 | 25939 | 1425 | 0.0521 |
| ObDBb5_aligned | 6150 | 6142 | 8 | 0.0013 | ObDBb5_aligned | 27364 | 25388 | 1976 | 0.0722 |
| ObOFa1_aligned | 6150 | 6149 | 1 | 0.0002 | ObOFa1_aligned | 27364 | 25980 | 1384 | 0.0506 |
| ObOFa2_aligned | 6150 | 6149 | 1 | 0.0002 | ObOFa2_aligned | 27364 | 24904 | 2460 | 0.0899 |
| ObOFa3_aligned | 6150 | 6150 | 0 | 0 | ObOFa3_aligned | 27364 | 25982 | 1382 | 0.0505 |
| ObOFa4_aligned | 6150 | 6149 | 1 | 0.0002 | ObOFa4_aligned | 27364 | 25875 | 1489 | 0.0544 |
| ObOFa5_aligned | 6150 | 6147 | 3 | 0.0005 | ObOFa5_aligned | 27364 | 26172 | 1192 | 0.0436 |
| ObAFa1_aligned | 6150 | 6149 | 1 | 0.0002 | ObAFa1_aligned | 27364 | 26720 | 644 | 0.0235 |
| ObAFa2_aligned | 6150 | 6147 | 3 | 0.0005 | ObAFa2_aligned | 27364 | 26602 | 762 | 0.0278 |
| ObAFa3_aligned | 6150 | 6146 | 4 | 0.0007 | ObAFa3_aligned | 27364 | 24369 | 2995 | 0.1095 |
| ObAFa4_aligned | 6150 | 6147 | 3 | 0.0005 | ObAFa4_aligned | 27364 | 25524 | 1840 | 0.0672 |
| ObAFa5_aligned | 6150 | 6147 | 3 | 0.0005 | ObAFa5_aligned | 27364 | 26200 | 1164 | 0.0425 |
| ObBFa1_aligned | 6150 | 6150 | 0 | 0 | ObBFa1_aligned | 27364 | 25834 | 1530 | 0.0559 |
| ObBFa2_aligned | 6150 | 6148 | 2 | 0.0003 | ObBFa2_aligned | 27364 | 25875 | 1489 | 0.0544 |
| ObBFa3_aligned | 6150 | 6150 | 0 | 0 | ObBFa3_aligned | 27364 | 27236 | 128 | 0.0047 |
| ObBFa4_aligned | 6150 | 6150 | 0 | 0 | ObBFa4_aligned | 27364 | 27304 | 60 | 0.0022 |
| ObBFa5_aligned | 6150 | 6149 | 1 | 0.0002 | ObBFa5_aligned | 27364 | 25932 | 1432 | 0.0523 |
| ObBFb1_aligned | 6150 | 6148 | 2 | 0.0003 | ObBFb1_aligned | 27364 | 22539 | 4825 | 0.1763 |
| ObBFb2_aligned | 6150 | 6150 | 0 | 0 | ObBFb2_aligned | 27364 | 27291 | 73 | 0.0027 |
| ObBFb3_aligned | 6150 | 6150 | 0 | 0 | ObBFb3_aligned | 27364 | 27236 | 128 | 0.0047 |
| ObBFb4_aligned | 6150 | 6150 | 0 | 0 | ObBFb4_aligned | 27364 | 26918 | 446 | 0.0163 |
| ObBFb5_aligned | 6150 | 6150 | 0 | 0 | ObBFb5_aligned | 27364 | 25960 | 1404 | 0.0513 |
| ObCMa1_aligned | 6150 | 6148 | 2 | 0.0003 | ObCMa1_aligned | 27364 | 27060 | 304 | 0.0111 |
| ObCMa2_aligned | 6150 | 6150 | 0 | 0 | ObCMa2_aligned | 27364 | 27227 | 137 | 0.005 |
| ObCMa3_aligned | 6150 | 6147 | 3 | 0.0005 | ObCMa3_aligned | 27364 | 27087 | 277 | 0.0101 |
| ObCMa4_aligned | 6150 | 6150 | 0 | 0 | ObCMa4_aligned | 27364 | 27060 | 304 | 0.0111 |
| ObHMa1_aligned | 6150 | 6150 | 0 | 0 | ObHMa1_aligned | 27364 | 27012 | 352 | 0.0129 |
| ObHMa2_aligned | 6150 | 6149 | 1 | 0.0002 | ObHMa2_aligned | 27364 | 26935 | 429 | 0.0157 |
| ObHMa3_aligned | 6150 | 6150 | 0 | 0 | ObHMa3_aligned | 27364 | 26972 | 392 | 0.0143 |
| ObHMa4_aligned | 6150 | 6149 | 1 | 0.0002 | ObHMa4_aligned | 27364 | 26956 | 408 | 0.0149 |
| ObHMa5_aligned | 6150 | 6148 | 2 | 0.0003 | ObHMa5_aligned | 27364 | 26981 | 383 | 0.014 |
| ObHMb1_aligned | 6150 | 6149 | 1 | 0.0002 | ObHMb1_aligned | 27364 | 27066 | 298 | 0.0109 |
| ObHMb2_aligned | 6150 | 6146 | 4 | 0.0007 | ObHMb2_aligned | 27364 | 27110 | 254 | 0.0093 |
| ObHMb3_aligned | 6150 | 6139 | 11 | 0.0018 | ObHMb3_aligned | 27364 | 26919 | 445 | 0.0163 |
| ObHMb4_aligned | 6150 | 6148 | 2 | 0.0003 | ObHMb4_aligned | 27364 | 27048 | 316 | 0.0115 |
| ObHMb5_aligned | 6150 | 6150 | 0 | 0 | ObHMb5_aligned | 27364 | 27127 | 237 | 0.0087 |

Denovo
| sample | n_loci | present_loci | missing_loci | frequency_missing | sample | n_sites | present_sites | missing_sites | frequency_missing |
| --- | --- | --- | --- | --- | --- | --- | --- | --- | --- |
| ObBDa1 | **4405** | 3489 | 916 | 0.2079 | ObBDa1 | **9053** | 7601 | 1452 | 0.1604 |
| ObSFa2 | **4405** | 3058 | 1347 | 0.3058 | ObSFa2 | **9053** | 7020 | 2033 | 0.2246 |
| ObSFa3 | **4405** | 3029 | 1376 | 0.3124 | ObSFa3 | **9053** | 6989 | 2064 | 0.228 |
| ObSFa4 | **4405** | 3839 | 566 | 0.1285 | ObSFa4 | **9053** | 7688 | 1365 | 0.1508 |
| ObSFa5 | **4405** | 4038 | 367 | 0.0833 | ObSFa5 | **9053** | 7796 | 1257 | 0.1388 |
| ObDBa1 | **4405** | 4138 | 267 | 0.0606 | ObDBa1 | **9053** | 8003 | 1050 | 0.116 |
| ObDBa2 | **4405** | 4142 | 263 | 0.0597 | ObDBa2 | **9053** | 8041 | 1012 | 0.1118 |
| ObDBa3 | **4405** | 4186 | 219 | 0.0497 | ObDBa3 | **9053** | 8310 | 743 | 0.0821 |
| ObDBa4 | **4405** | 4116 | 289 | 0.0656 | ObDBa4 | **9053** | 8035 | 1018 | 0.1124 |
| ObDBa5 | **4405** | 4255 | 150 | 0.0341 | ObDBa5 | **9053** | 8100 | 953 | 0.1053 |
| ObDBb1 | **4405** | 4182 | 223 | 0.0506 | ObDBb1 | **9053** | 8078 | 975 | 0.1077 |
| ObDBb2 | **4405** | 4294 | 111 | 0.0252 | ObDBb2 | **9053** | 8531 | 522 | 0.0577 |
| ObDBb3 | **4405** | 4310 | 95 | 0.0216 | ObDBb3 | **9053** | 8499 | 554 | 0.0612 |
| ObDBb4 | **4405** | 4203 | 202 | 0.0459 | ObDBb4 | **9053** | 8462 | 591 | 0.0653 |
| ObDBb5 | **4405** | 4147 | 258 | 0.0586 | ObDBb5 | **9053** | 8189 | 864 | 0.0954 |
| ObOFa1 | **4405** | 4239 | 166 | 0.0377 | ObOFa1 | **9053** | 8522 | 531 | 0.0587 |
| ObOFa2 | **4405** | 3949 | 456 | 0.1035 | ObOFa2 | **9053** | 7673 | 1380 | 0.1524 |
| ObOFa3 | **4405** | 4266 | 139 | 0.0316 | ObOFa3 | **9053** | 8421 | 632 | 0.0698 |
| ObOFa4 | **4405** | 4322 | 83 | 0.0188 | ObOFa4 | **9053** | 8488 | 565 | 0.0624 |
| ObOFa5 | **4405** | 4221 | 184 | 0.0418 | ObOFa5 | **9053** | 8544 | 509 | 0.0562 |
| ObAFa1 | **4405** | 4252 | 153 | 0.0347 | ObAFa1 | **9053** | 8697 | 356 | 0.0393 |
| ObAFa2 | **4405** | 4236 | 169 | 0.0384 | ObAFa2 | **9053** | 8644 | 409 | 0.0452 |
| ObAFa3 | **4405** | 3831 | 574 | 0.1303 | ObAFa3 | **9053** | 7623 | 1430 | 0.158 |
| ObAFa4 | **4405** | 3981 | 424 | 0.0963 | ObAFa4 | **9053** | 8053 | 1000 | 0.1105 |
| ObAFa5 | **4405** | 4112 | 293 | 0.0665 | ObAFa5 | **9053** | 8398 | 655 | 0.0724 |
| ObBFa1 | **4405** | 4138 | 267 | 0.0606 | ObBFa1 | **9053** | 8236 | 817 | 0.0902 |
| ObBFa2 | **4405** | 4097 | 308 | 0.0699 | ObBFa2 | **9053** | 8348 | 705 | 0.0779 |
| ObBFa3 | **4405** | 4358 | 47 | 0.0107 | ObBFa3 | **9053** | 8959 | 94 | 0.0104 |
| ObBFa4 | 4405 | 4363 | 42 | 0.0095 | ObBFa4 | 9053 | 8987 | 66 | 0.0073 |
| ObBFa5 | 4405 | 4087 | 318 | 0.0722 | ObBFa5 | 9053 | 8303 | 750 | 0.0828 |
| ObBFb1 | 4405 | 3443 | 962 | 0.2184 | ObBFb1 | 9053 | 6802 | 2251 | 0.2486 |
| ObBFb2 | 4405 | 4366 | 39 | 0.0089 | ObBFb2 | 9053 | 8969 | 84 | 0.0093 |
| ObBFb3 | 4405 | 4363 | 42 | 0.0095 | ObBFb3 | 9053 | 8953 | 100 | 0.011 |
| ObBFb4 | 4405 | 4311 | 94 | 0.0213 | ObBFb4 | 9053 | 8799 | 254 | 0.0281 |
| ObBFb5 | 4405 | 4094 | 311 | 0.0706 | ObBFb5 | 9053 | 8262 | 791 | 0.0874 |
| ObCMa1 | 4405 | 4311 | 94 | 0.0213 | ObCMa1 | 9053 | 8757 | 296 | 0.0327 |
| ObCMa2 | 4405 | 4308 | 97 | 0.022 | ObCMa2 | 9053 | 8745 | 308 | 0.034 |
| ObCMa3 | 4405 | 4332 | 73 | 0.0166 | ObCMa3 | 9053 | 8746 | 307 | 0.0339 |
| ObCMa4 | 4405 | 4275 | 130 | 0.0295 | ObCMa4 | 9053 | 8736 | 317 | 0.035 |
| ObHMa1 | 4405 | 4360 | 45 | 0.0102 | ObHMa1 | 9053 | 8808 | 245 | 0.0271 |
| ObHMa2 | 4405 | 4336 | 69 | 0.0157 | ObHMa2 | 9053 | 8724 | 329 | 0.0363 |
| ObHMa3 | 4405 | 4308 | 97 | 0.022 | ObHMa3 | 9053 | 8663 | 390 | 0.0431 |
| ObHMa4 | 4405 | 4326 | 79 | 0.0179 | ObHMa4 | 9053 | 8766 | 287 | 0.0317 |
| ObHMa5 | 4405 | 4297 | 108 | 0.0245 | ObHMa5 | 9053 | 8832 | 221 | 0.0244 |
| ObHMb1 | 4405 | 4335 | 70 | 0.0159 | ObHMb1 | 9053 | 8833 | 220 | 0.0243 |
| ObHMb2 | 4405 | 4340 | 65 | 0.0148 | ObHMb2 | 9053 | 8882 | 171 | 0.0189 |
| ObHMb3 | 4405 | 4284 | 121 | 0.0275 | ObHMb3 | 9053 | 8708 | 345 | 0.0381 |
| ObHMb4 | 4405 | 4347 | 58 | 0.0132 | ObHMb4 | 9053 | 8865 | 188 | 0.0208 |
| ObHMb5 | 4405 | 4360 | 45 | 0.0102 | ObHMb5 | 9053 | 8861 | 192 | 0.0212 |

